# Tree Surface Area Allometry

**DOI:** 10.1101/2024.04.23.590783

**Authors:** Alexander Shenkin, Phil Wilkes, Brian Enquist, Alvaro Sarmiento, Andy Burt, Kim Calders, Pasi Raumonen, Lisa Patrick Bentley, Mat Disney, Yadvinder Malhi

## Abstract

Utilizing terrestrial laser scanning (TLS) and three-dimensional modeling, this study quantitatively assessed the woody surface areas of 2161 trees across ecosystems encompassing both tropical and temperate forests. TLS enables precise measurement of tree structures at unprecedented scales. This research builds on theoretical scaling relationships with empirical data, significantly refining our understanding of tree woody surface area. Key findings indicate that direct measurements diverge from theoretical predictions, particularly in the finer branch structures, suggesting modifications to existing allometric models might be necessary. This integration of direct measurements with TLS not only challenges established theories but also enhances our capability to accurately model tree surface area, which is key for understanding forest carbon dynamics and metabolic scaling in ecological systems.

## Introduction

Biological surface areas constrain fluxes of material and energy. The woody surface area of a tree is an important quantity that constrains respiration, the fluxes of gases such as CO2 and CH4, radiative energy, and water and nutrient budgets (Whittaker and Woodwell 1967). It is particularly important in the scaling of woody respiration, which itself is an important and poorly constrained component of the forest carbon cycle. West *et al*. (West, Brown, and Enquist 1999) suggest that organisms have evolved to maximize their metabolic rates, and these metabolic rates are in part controlled by the surface area across which organisms exchange material and energy with the environment. Increasing surface areas must be balanced against other trade-offs such as mechanical stability and the efficiency of resource transportation within the organism.

The relationship between the size of an organism and the rate of its metabolic processes, which relate to fluxes across the surface area of its body, is a core concept of biology (Kleiber 1932).

Measuring surface areas of large trees is difficult, and traditionally requires climbing and measuring trees or felling them and measuring them on the ground. A number of efforts have used Recent advances in terrestrial laser scanning (TLS) technology and 3D modeling algorithms have made it possible to measure the woody structure of many trees to very fine scales non destructively. Thus, the scales across which we are able to directly measure woody structure have expanded scales of magnitude both upwards to encompass more trees, and downwards to encompass finer branching. Here we take advantage of these advances to quantify the woody surface areas of 2161 trees from 12 plots across tropical and temperate ecosystems, environmental gradients, evolutionary histories, and biogeographic regions.

Even when felled, measuring the finer branches of trees is difficult and time consuming. Thus, many previous efforts to understand woody surface area have relied on theoretical scaling relationships to make predictions of the amount of smaller branch material in a tree (e.g. Chambers et al. 2004; Whittaker and Woodwell 1967). Here we couple TLS’s ability to directly measure these small structures with scans of felled branches to quantify the surface area held by these small structures, and evaluate enables us to directly measure these finer structures to a certain level. We make theoretical predictions of how surface area is distributed across branch size classes, and we combine TLS of whole trees with focused scanning of branches to evaluate those predictions in trees and across entire plots.

### Current allometries

While a number of empirical relationships between tree characteristics (e.g. DBH, height) and woody surface areas have been developed (e.g. Weiskittel and Maguire 2006; Halldin 1985), two models provide theoretical predictions for the woody surface area of a tree: Pipe Model Theory (Shinozaki et al. 1964), and Metabolic Scaling Theory (West, Brown, and Enquist 1997).

### Pipe Model Theory

Shinozaki’s (1964) pipe model theory posits that the diameter frequency of branches in a tree (*ϕ*(*d*)) decreases as a power of the branch diameter *d*:

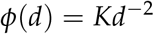

Shinozaki (1964) and others have shown that the power relationship varies between -1.5 – -2, though most subsequent authors assume the exponenent to be -2. Primary stems are modeleled apart from branches as cones, as:

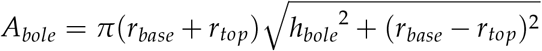

where *A*_*bole*_ is the surface area of the bole, *r*_*base*_ and *r*_*top*_ are the radii at the base and top of the bole respectively, and *h*_*bole*_ is the height of the bole.

Yoneda (1993) defines the constant *K* as *K* = 4*m*_*b*_/(*πρ*(*d*_*max*_/*d*_*min*_)), where *m*_*b*_ is the dry mass of branches, *ρ* is bulk density, and *d*_*max*_ and *d*_*min*_ are the maximum and minimum branch diameters. In practice, authors make different assumptions of branch diameter limits: estimates of *d*_*max*_ vary between the size of the trunk at the base of the first fork (*d*_*b*_), and 0.6*d*_*b*_; *d*_*min*_ is often assumed to be 2mm.

Chambers (2004) employed Yoneda’s (1993) branch and stem formulae to 315 felled trees in the Amazon, directly measuring *r*_*base*_, *h*_*bole*_, and *m*_*b*_, to produce a whole-tree surface area allometry based solely on DBH (*D*_*base*_):

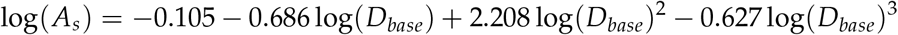

In most cases we measured branch tip widths, and

### Metabolic Scaling Theory

West et al. (1997) derives scaling relationships for organisms based on optimal resource transport efficiency. This optimality requires a fractal branching network, with successive branching generations related to the previous ones as *β* = *r*_*k*+1_/*r*_*k*_ and *γ* = *l*_*k*+1_/*l*_*k*_, where *k* is the branching order of which there are *N* orders. Thus, 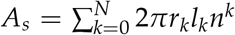, where *n* is the furcation ratio (usually 2). Hence, as a geometric series,

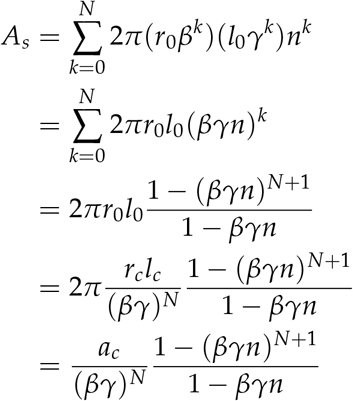

where 2*πr*_0_*l*_0_ is the surface area of the tree bole to the first branch. Alternatively, as *r*_*N*_ = *r*_0_*β*^*N*^ and *l*_*N*_ = *l*_0_*γ*^*N*^, and *r*_*c*_ *≡ r*_*N*_, *l*_*c*_ *≡ l*_*N*_, and the surface area of a “capillary” or terminal twig is *a*_*c*_ = 2*πr*_*c*_*l*_*c*_, 2*πr*_0_*l*_0_ can be rewritten in terms of the terminal twigs, as 2*πr*_*c*_*l*_*c*_(*βγ*)^*−N*^ = *a*_*c*_(*βγ*)^*−N*^.

This gives us an expression for *A*_*s*_ in terms of branching ratios and order numbers. We now turn to relationships between resource flow and capillary numbers, following West, Brown, and Enquist (1997), to relate this expression for *A*_*s*_ to mass scaling exponents.

From West, Brown, and Enquist (1997) eq. 2:

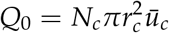

where *Q*_0_ is the volume rate of water flow and ū_*c*_ is the flow velocity in capillaries (or branch tips in this case). Thus, since capillaries are assumed invariant,

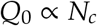

Metabolism (*B*) is posited to be directly related to the flow rate of resources, hence,

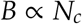

If *B* ∝ *M*^*a*^, where *M* is total mass of the organism and *a* is shown to be 3/4, then

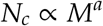

If surface area is related to mass such that *A*_*s*_^*z*^ ∝ *M*, then

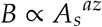

where *a* = 3/4. Thus, *Q*_0_ ∝ *A*_*s*_^*az*^, *N*_*c*_ ∝ *A*_*s*_^*az*^, and following West, Brown, and Enquist (1997) eq. 3,

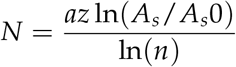

Substituting this into the previous equation relating *A*_*s*_ with branching ratios, we find

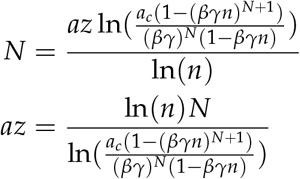

This is our expression for surface area scaling. We are unable to remove some terms that mass scaling does. Thus, we end up with a more complex expression that is a function of N.

Looking across N and setting *a*_*c*_ = 1, we find *a * z* = 0.52 for *N* = 1, and increases to an asymptote of *a * z* = 1.

**Figure 1.**
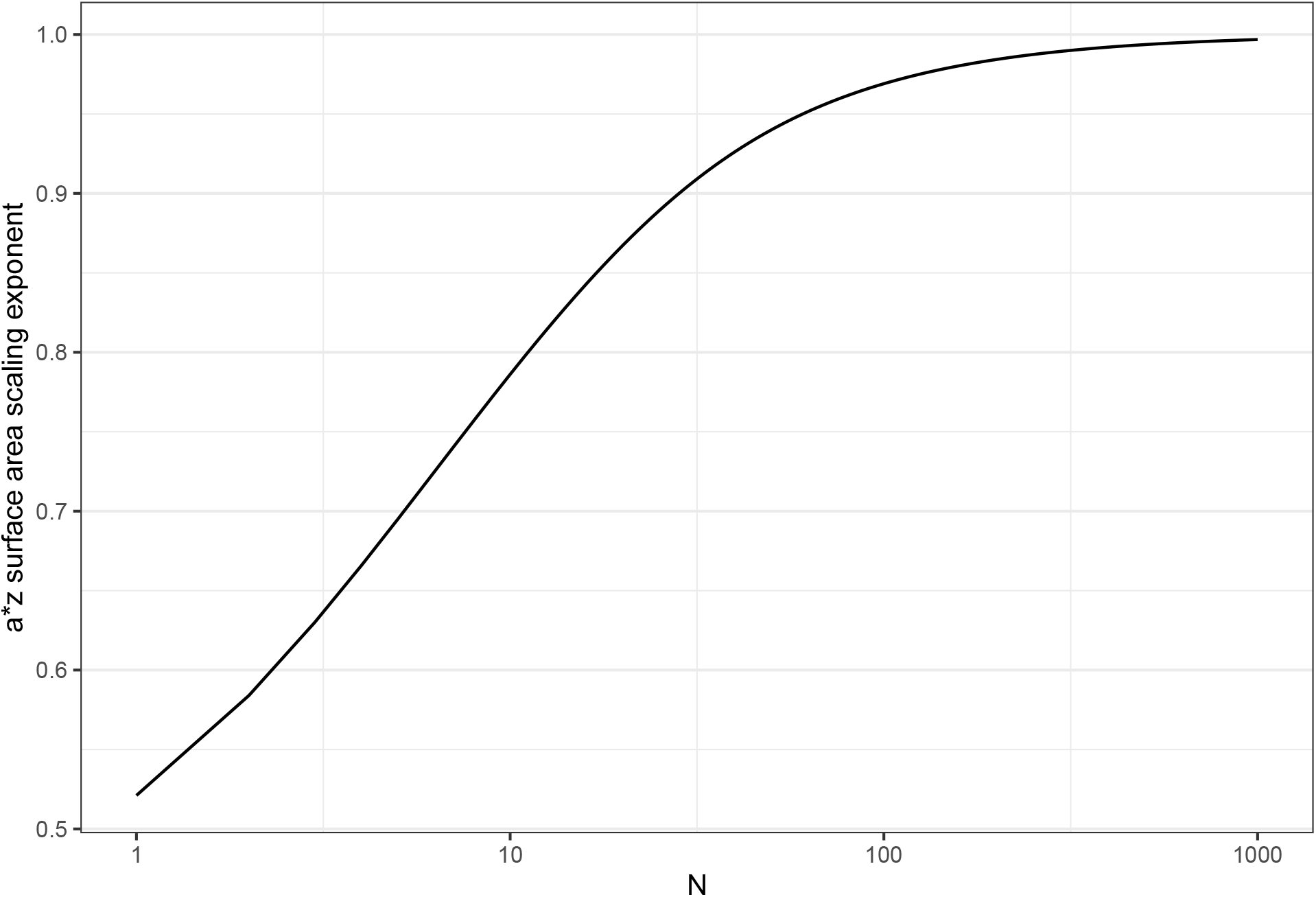
ST surface area scaling exponent (a * z) as a function of N, the number of branch orders in the tree

The scaling exponent relating surface area to mass is *z*: *M* ∝ *A*_*s*_^*z*^. According to MST, *a* = 3/4, thus, looking across N, we can examine the surface area - mass scaling exponent 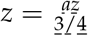. Here we find that *z* = 0.69 for *N* = 1, and increases to an asymptote of *z* = 1.33 = 4/3.

Because we cannot use the *nαβ <<* 1 assumption that works for volume scaling, we have to look at what N is likely to be as a function of DBH (or mass).

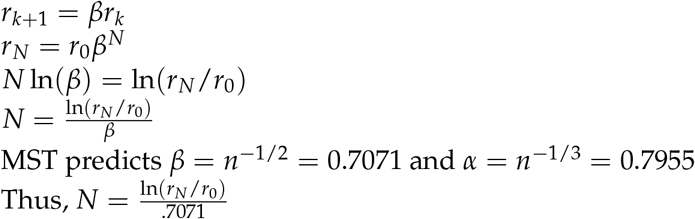

**Figure 2.**
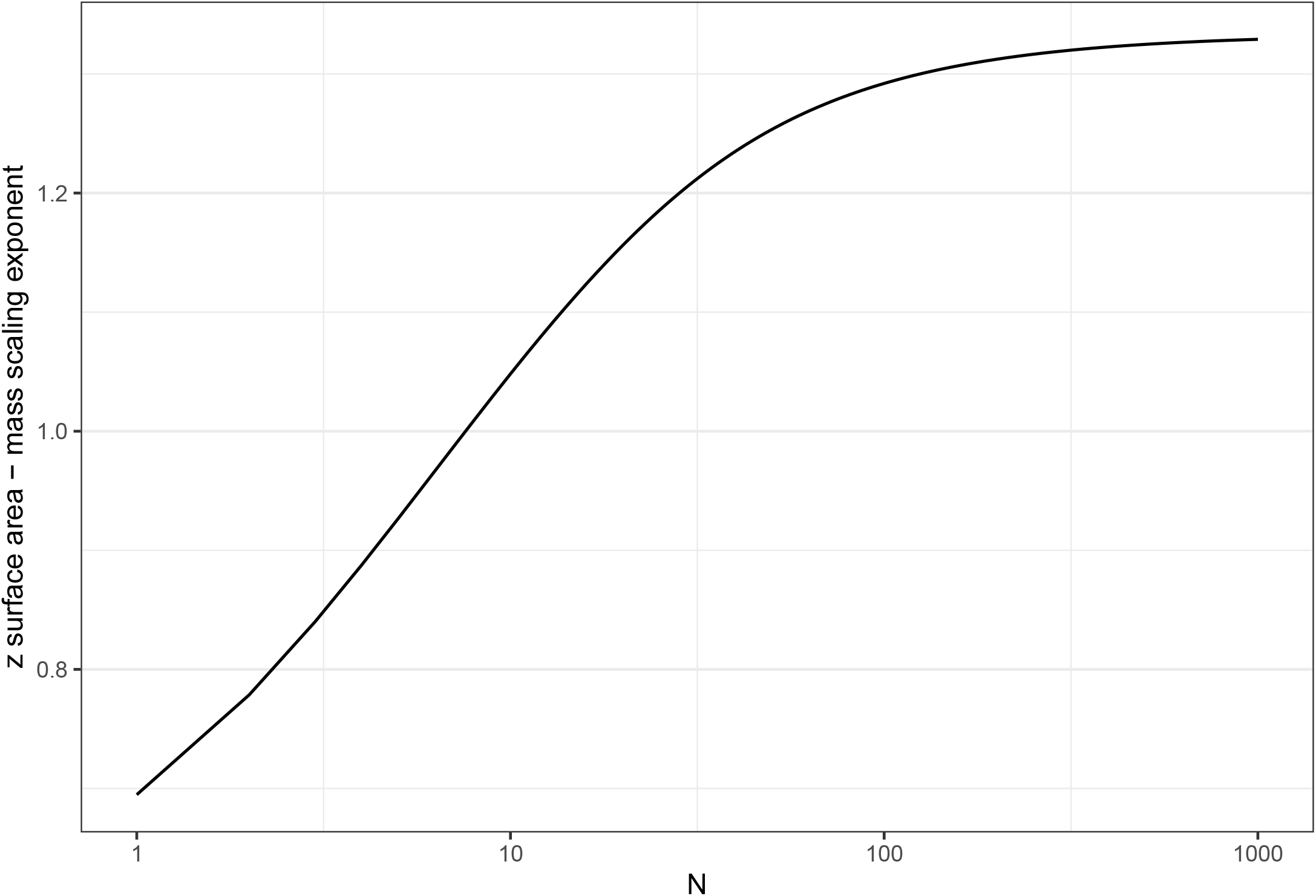
ST surface area - mass scaling exponent (z) as a function of N, the number of branch orders in the tree

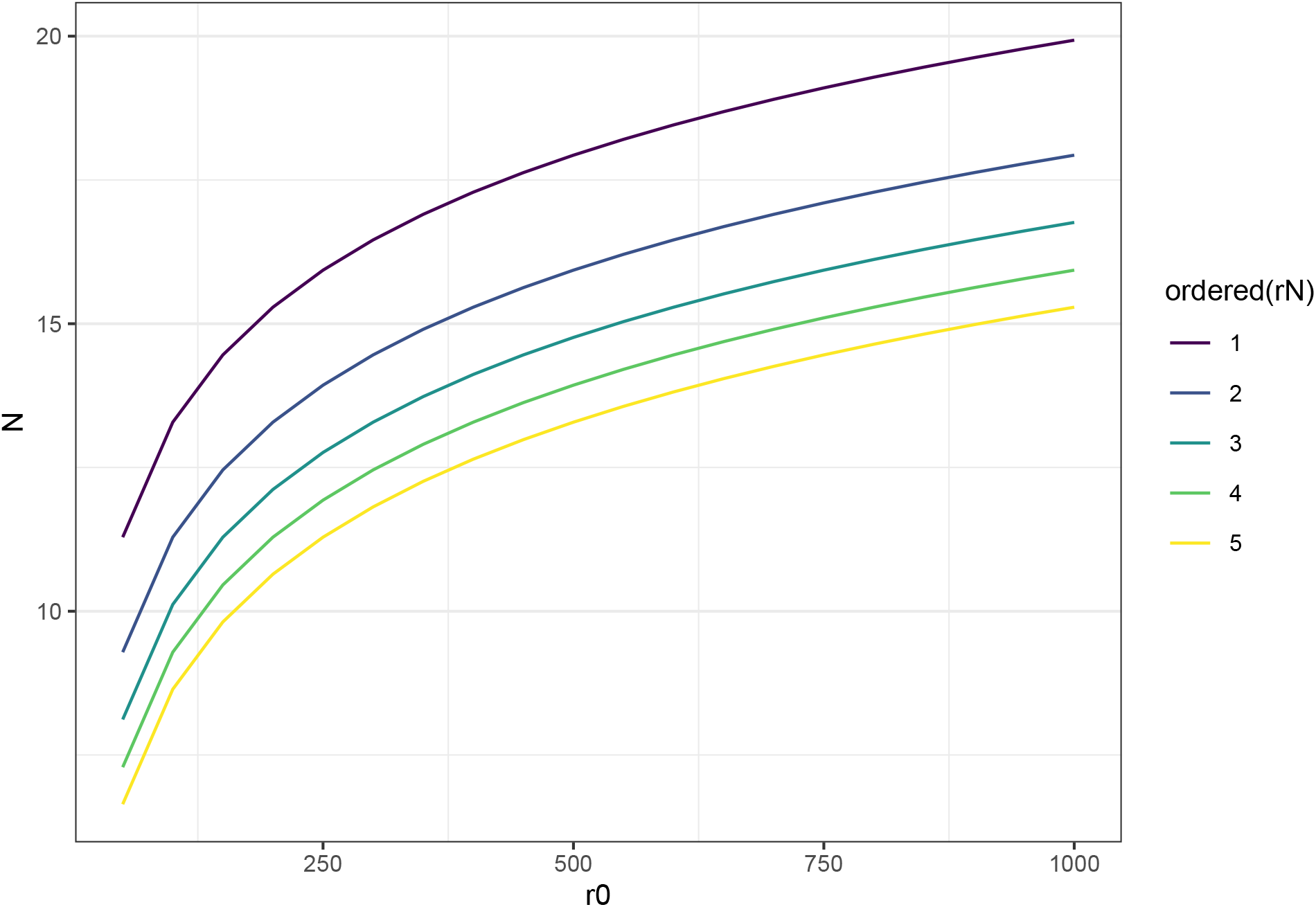

## Methods

### Sites

Validation conducted in Guyana, Peru, and Indonesia. Full-hectare analyses conducted in Wytham Woods, UK, and Cauxiana, Brazil.

### TLS operation

#### QSM & optimization

Pointcloud homogenization. Optimization: Treeqsm provides a metric that calculates the sum of distances from each point to the nearest cylinder.

QSMs are better off without leaf point removal, according to a small study by S. Moorthy (pers comm) of 5 trees in Wytham Woods scanned during summer (leaf on) and winter (leaf off).

**Table 1:**
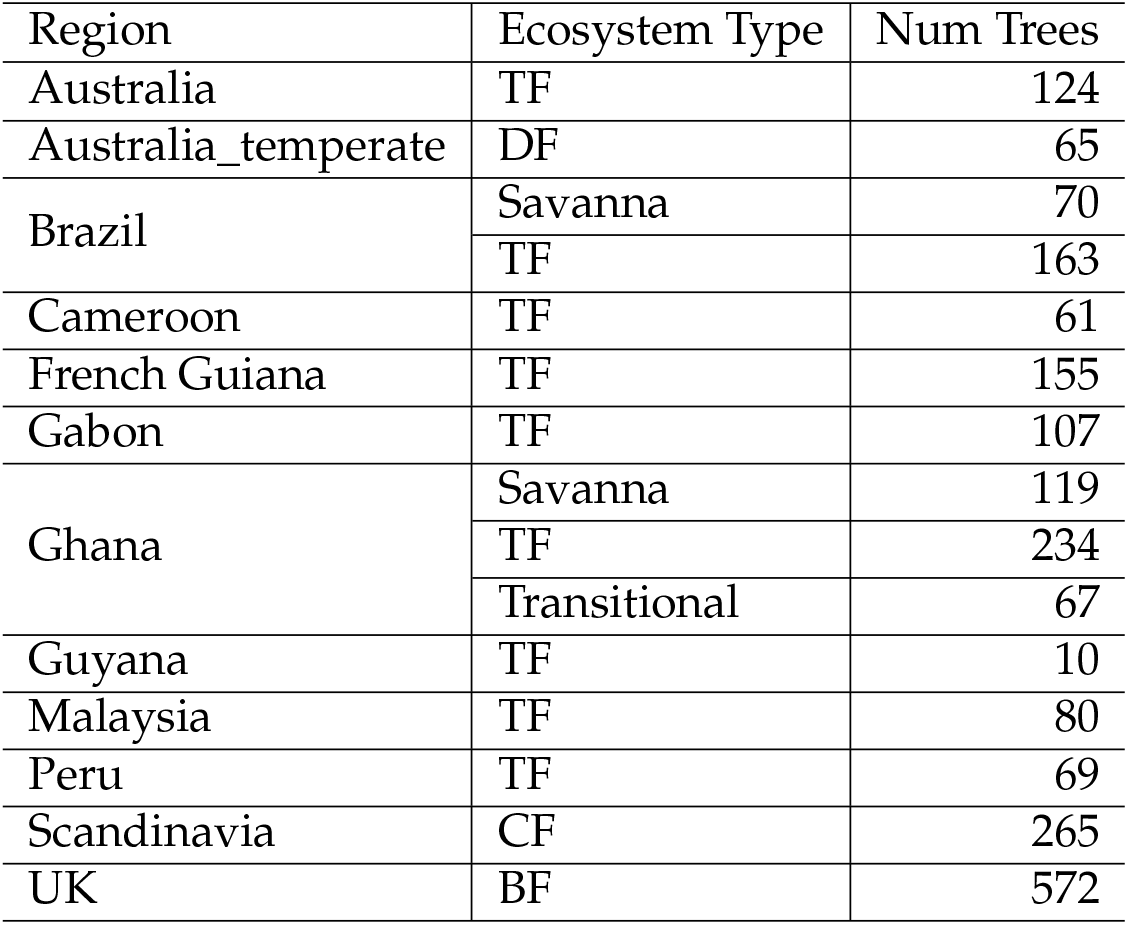
Number of trees per region in analysis.

## Results

### Validation

#### Branch validation

Here we validate the smaller branch diameter estimates using branch scan data.

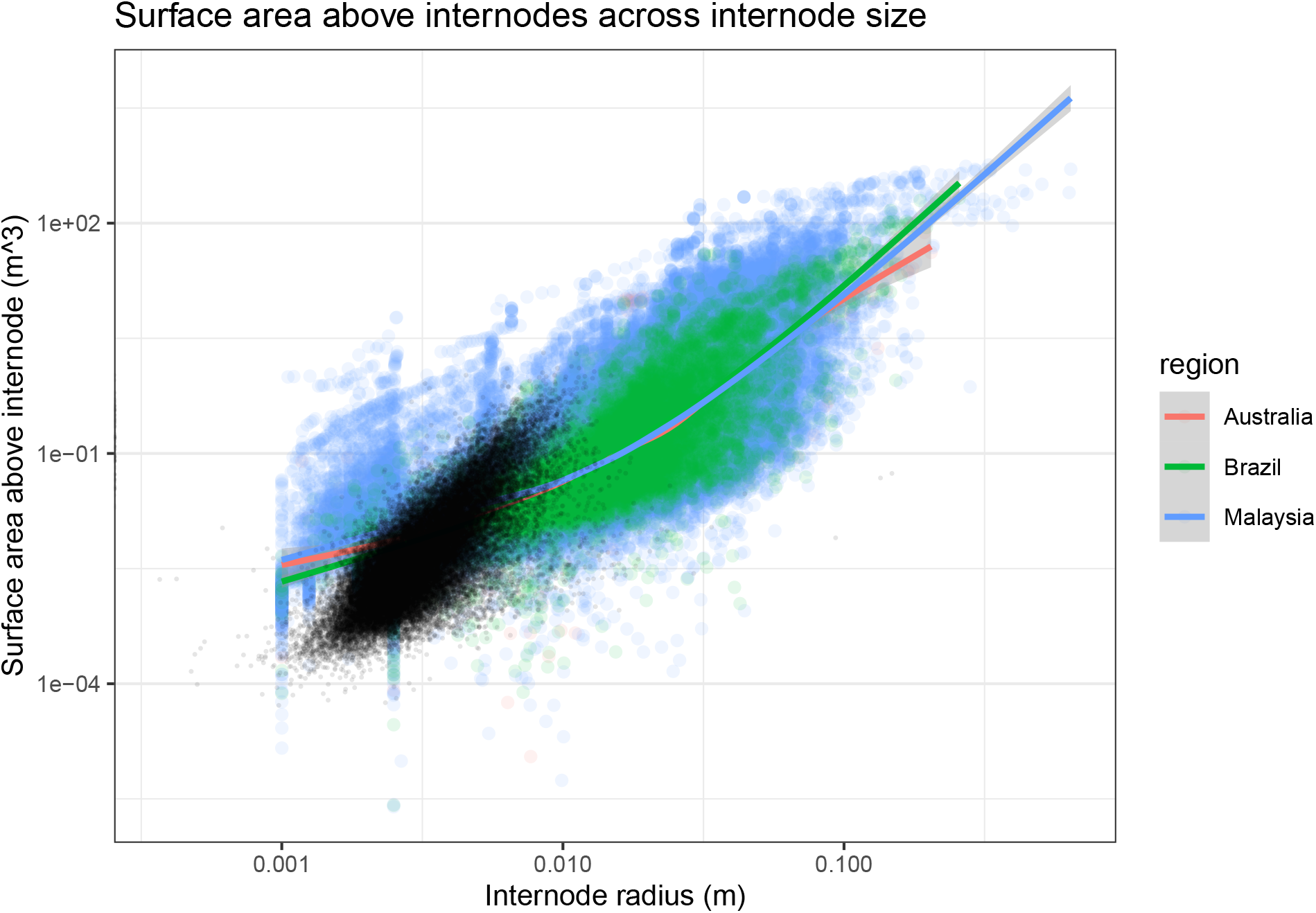

What seems to be the case here is that QSMs from forest-scanned trees are overestimating the amount of surface area above each internode in smaller branches by a factor of 10 or so. This much seems clear. Future work will test the *number* of branches of those size classes in the entire tree, versus the number that treeQSM thinks there are. A potential way of going about it would be to assume conservation of sapwood. Then, use the branches at, say, 10cm, and see how the total cross sectional area of sapwood goes down across, say, path distance from internode. We would expect some sort of variation, but if we get a drop in cross sectional area on the scale of an order of magnitude, then we might conclude that these effects are canceling each other out..

Let’s look at each plot. Take special note of NXV01 where we were scanning very close to the trees.

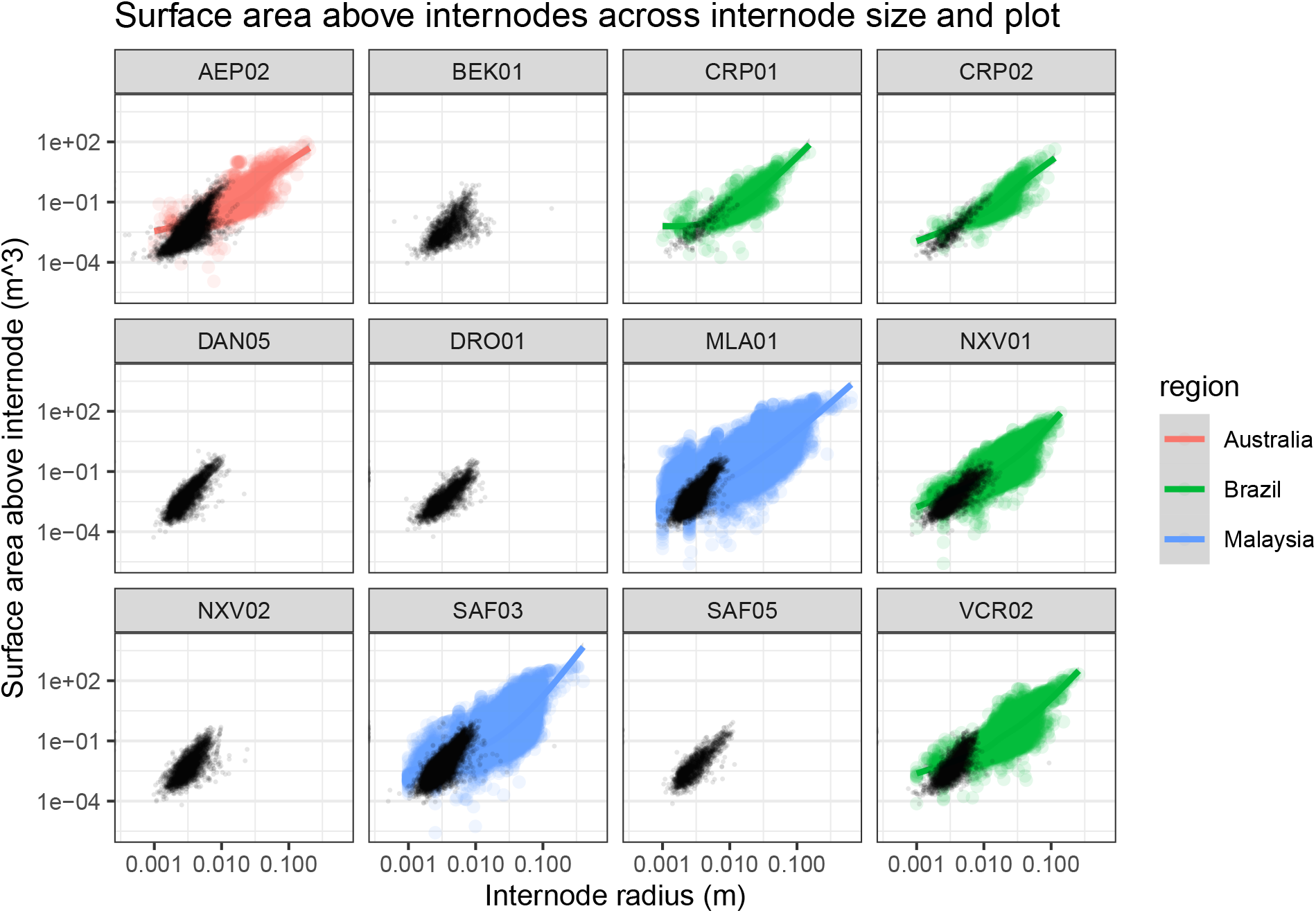

Validating each branch in the context of each tree, it actually looks like QSMs are *underestimating* the surface area above a branch, if anything.

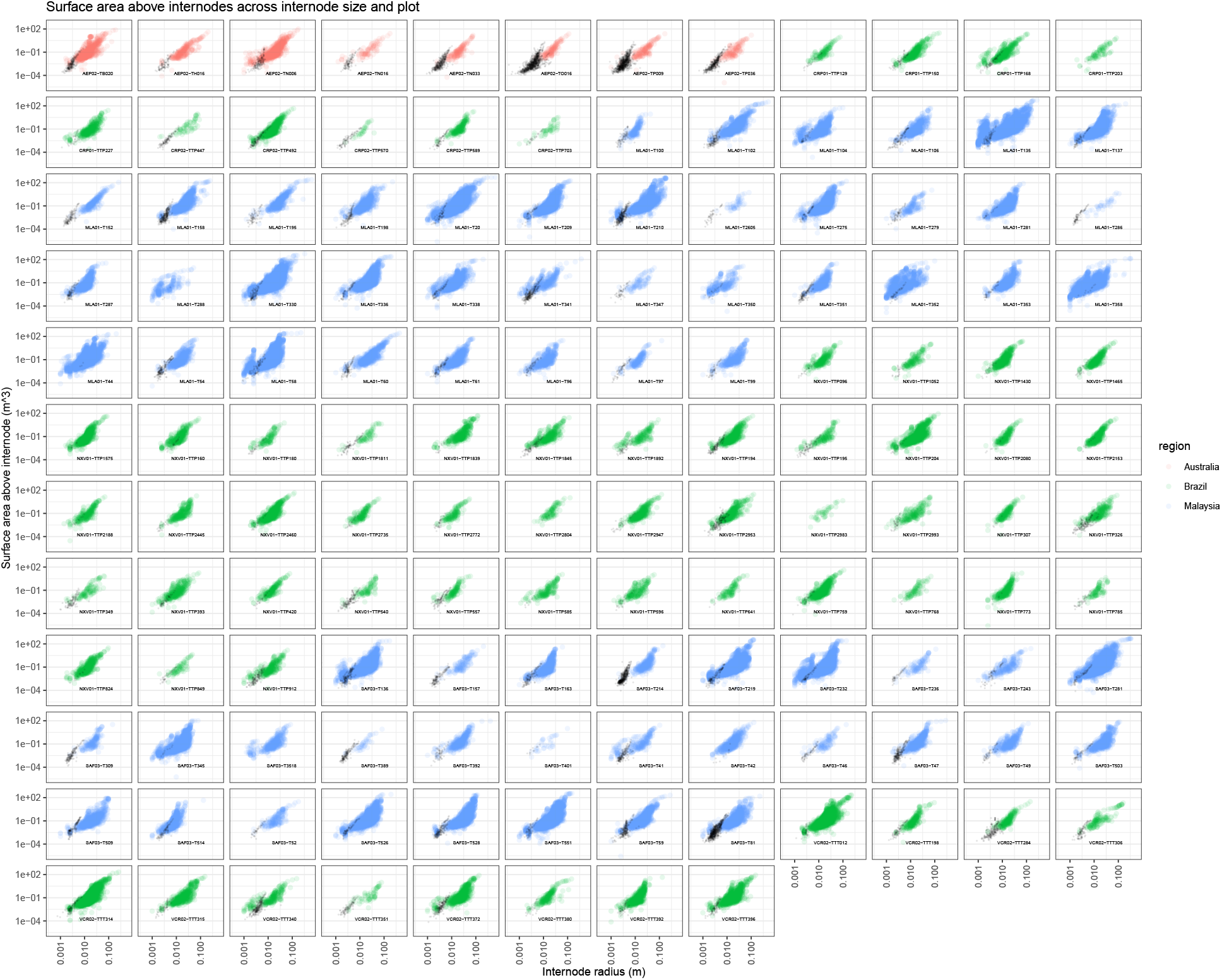

How do the closely-scanned branches compare to branches of similar sizes (+- 5mm radius) pruned from the QSM tree as scanned in the fields?

What we notice here: field QSM branches have lots of truncated members, it seems, from the many branches with small surface areas). Let’s look at the difference between the mean QSM branch surface area (aka predicted), and scanned branch surface ara (aka measured).

With very little predictive power evident in the measured vs. predicted figure, our main question then becomes, is there any bias?

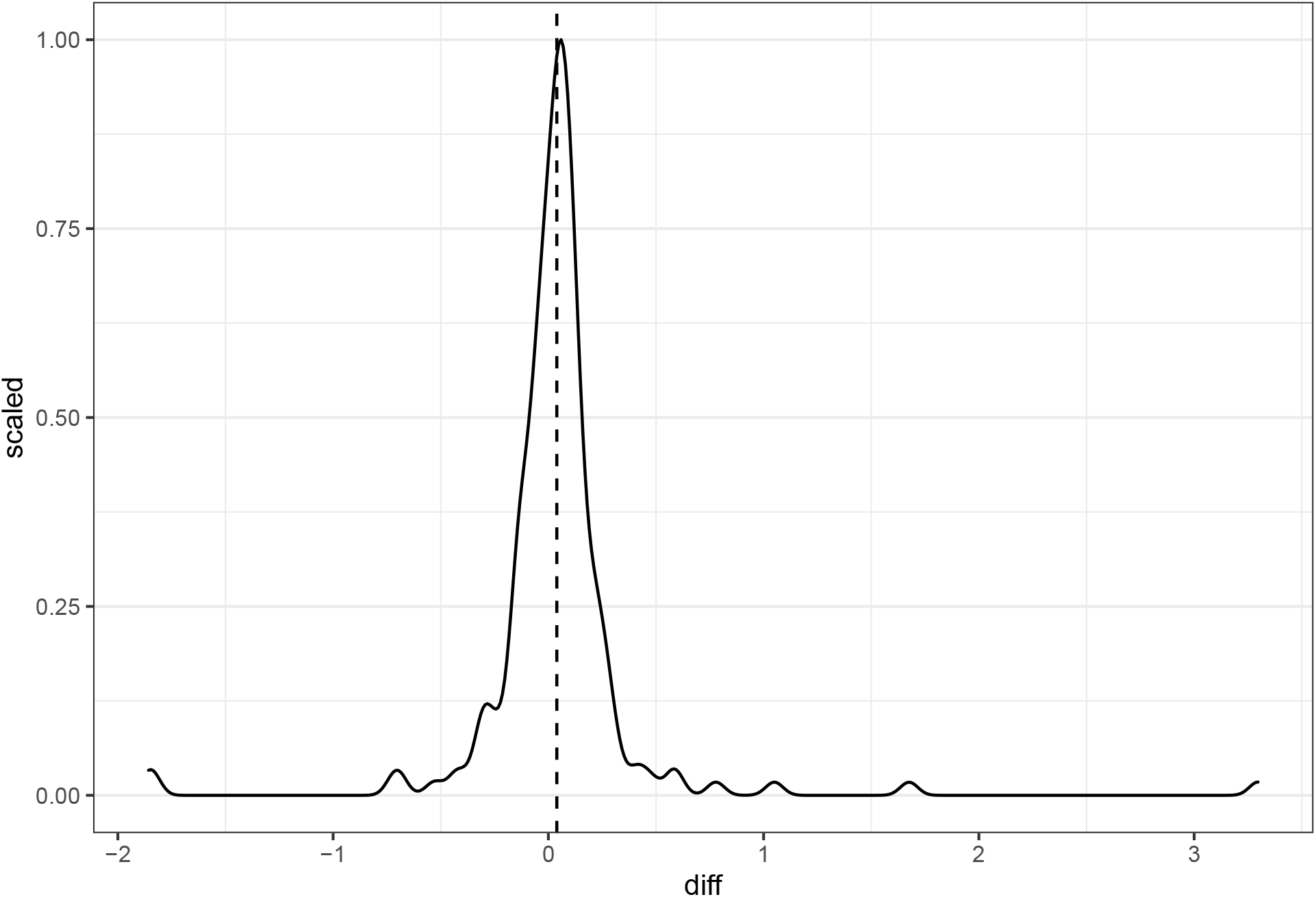

**Figure 3.**
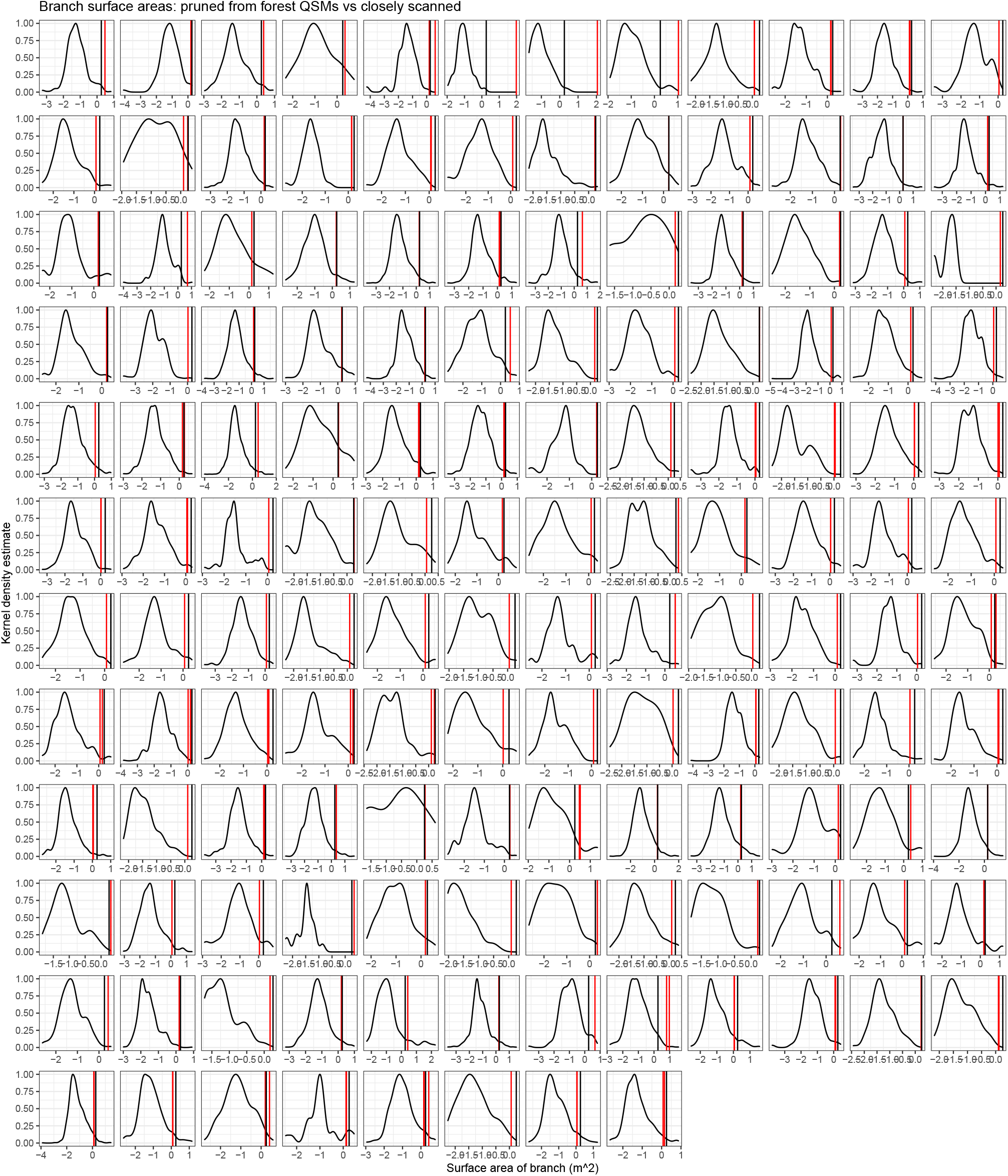
Kernel density estimates of surface areas of branches of similar basal diameter, their mean (vertical black line), and the surface area of the closely-scanned branch (vertical red line). Note that the x-axis is on a log scale for readability.

**Figure 4.**
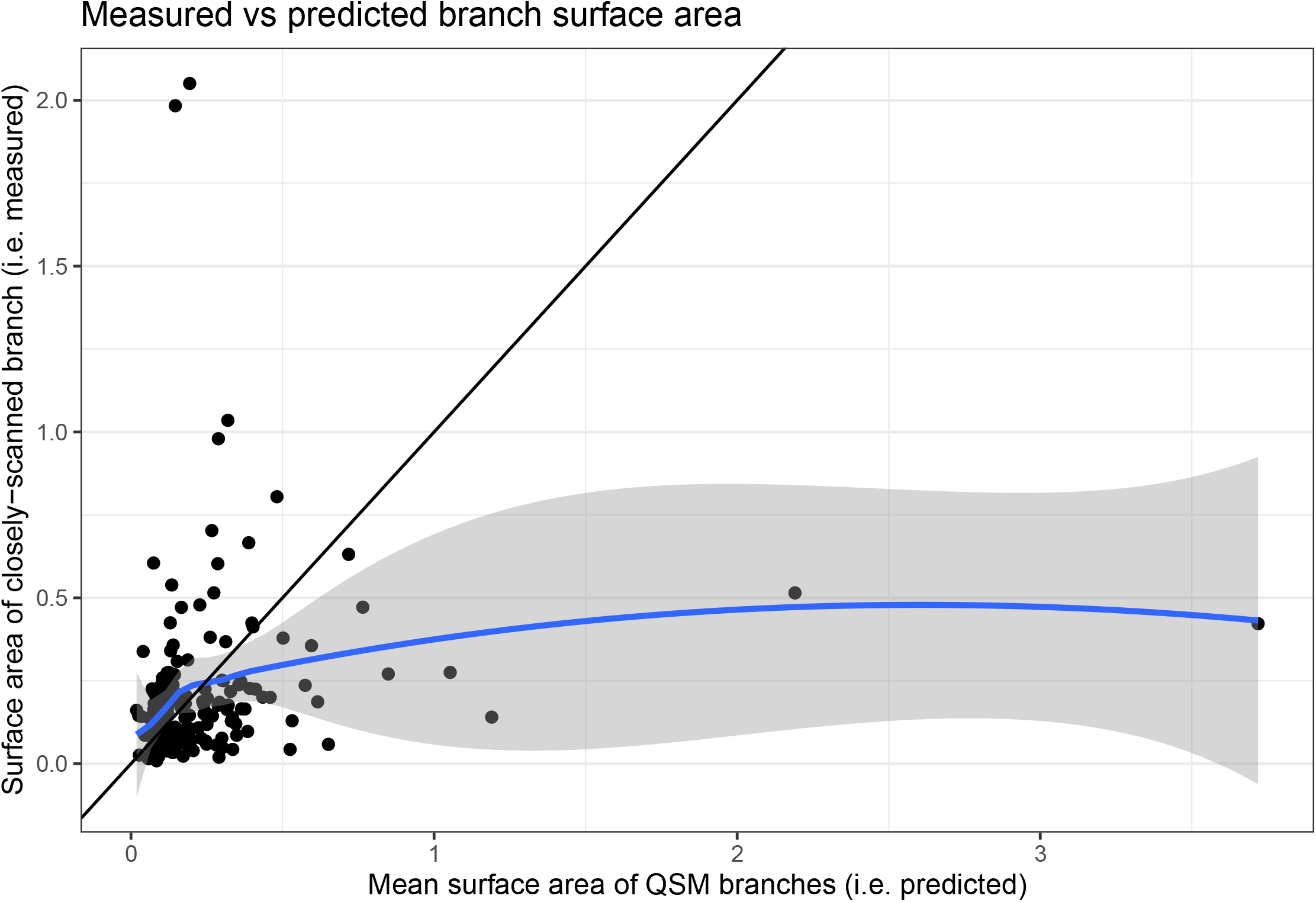
there is large variation here, and essentially no predictive power. Line is 1:1.

The mean difference between forest-QSM and closely-scanned branches is 0.0391595*m*^2^. Thus, while there is wide variation, forest-QSM branches have a slight per-branch bias of +4 *cm*^2^ in surface area compared to closely scanned branches, though a t-test indicates it is not a significant difference:

c(`mean` of x = 0.0391595337061303), c(t = 1.34813554552658), 0.179299497099954, c(df = 181), -0.0181551109201555, 0.096474178332416, One Sample t-test, two.sided

So, we conclude that, on a branch-by-branch basis, forest-QSM’s don’t pose a large risk for introducing bias. What we have not verified is that the *number* of small branches estimated with forest-scanned QSMs is not biased compared to the true number of branches. We hope to test that a later date; we don’t currently have the data to do it. Another way to approach the problem would be to make assumptions about conservation of cross-sectional area, and compare cross sectional area of the bole below first branch versus the cross-sectional area as one traverses out the branching network into smaller branches. We would expect the overall woody cross sectional area to remain more or less constant.

### Whole tree validation

In short, what we find is: removing leaf points is important. Once that is done, the entire non-truncated QSM should be used. This then is a quite conservative estimate of surface area. Further work could be developed to extrapolate using branch scans and scaling to estimate full surface areas.

### New surface area allometry

He were model per-tree surface area versus DBH across ecosystems.

### Data exploration - across plots

**Figure 5.**
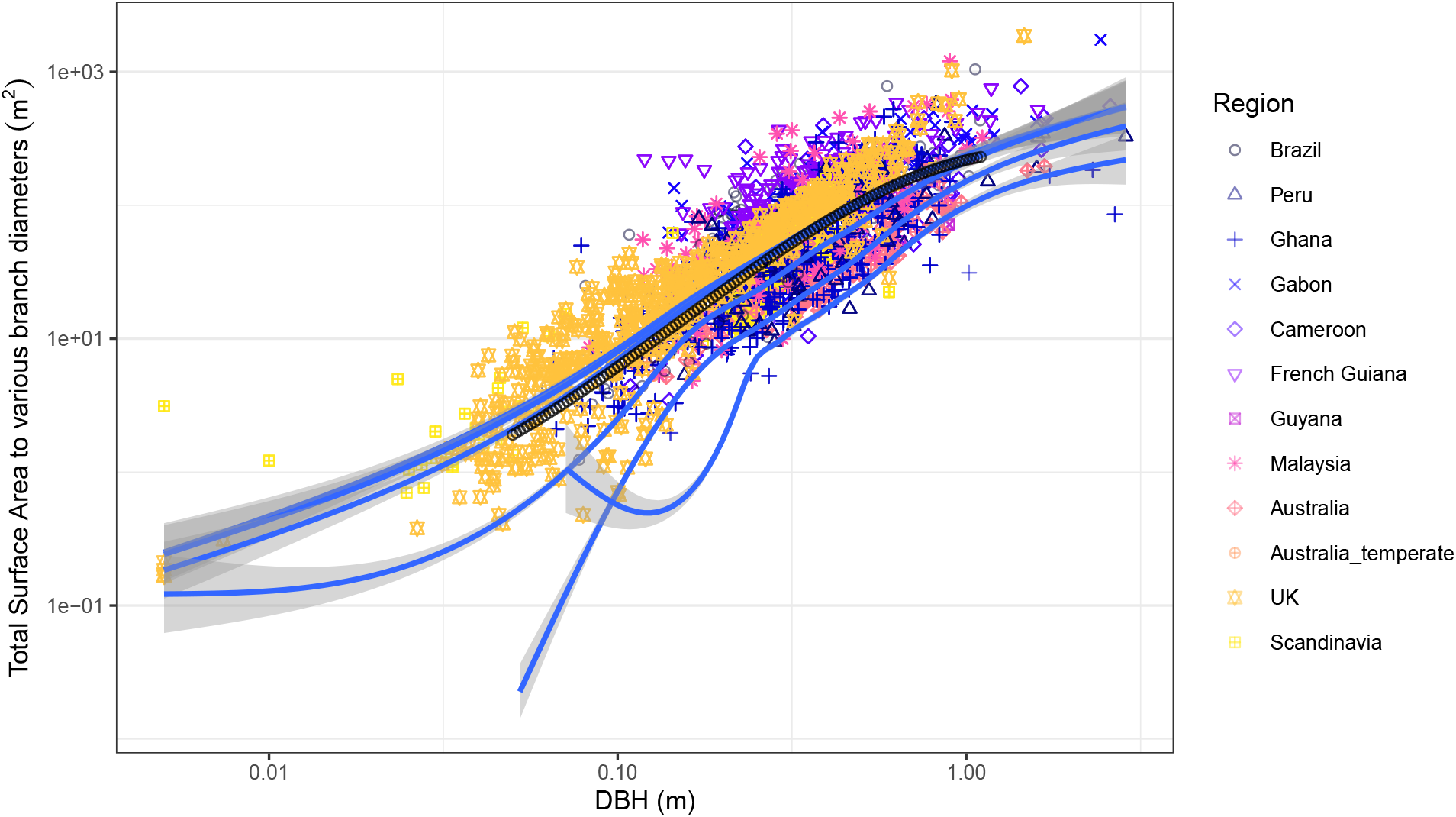
Surface area vs dbh across all trees. Each blue line includes increasingly fine branch diameters: 20cm (lowest line), 10cm, 5cm, 2cm, 1cm, and including all branches (top line). Black dots are expected values of Chambers (2004) allometry.

### Log-log scale

### Linear scale

### Linear model

**Figure 6.**
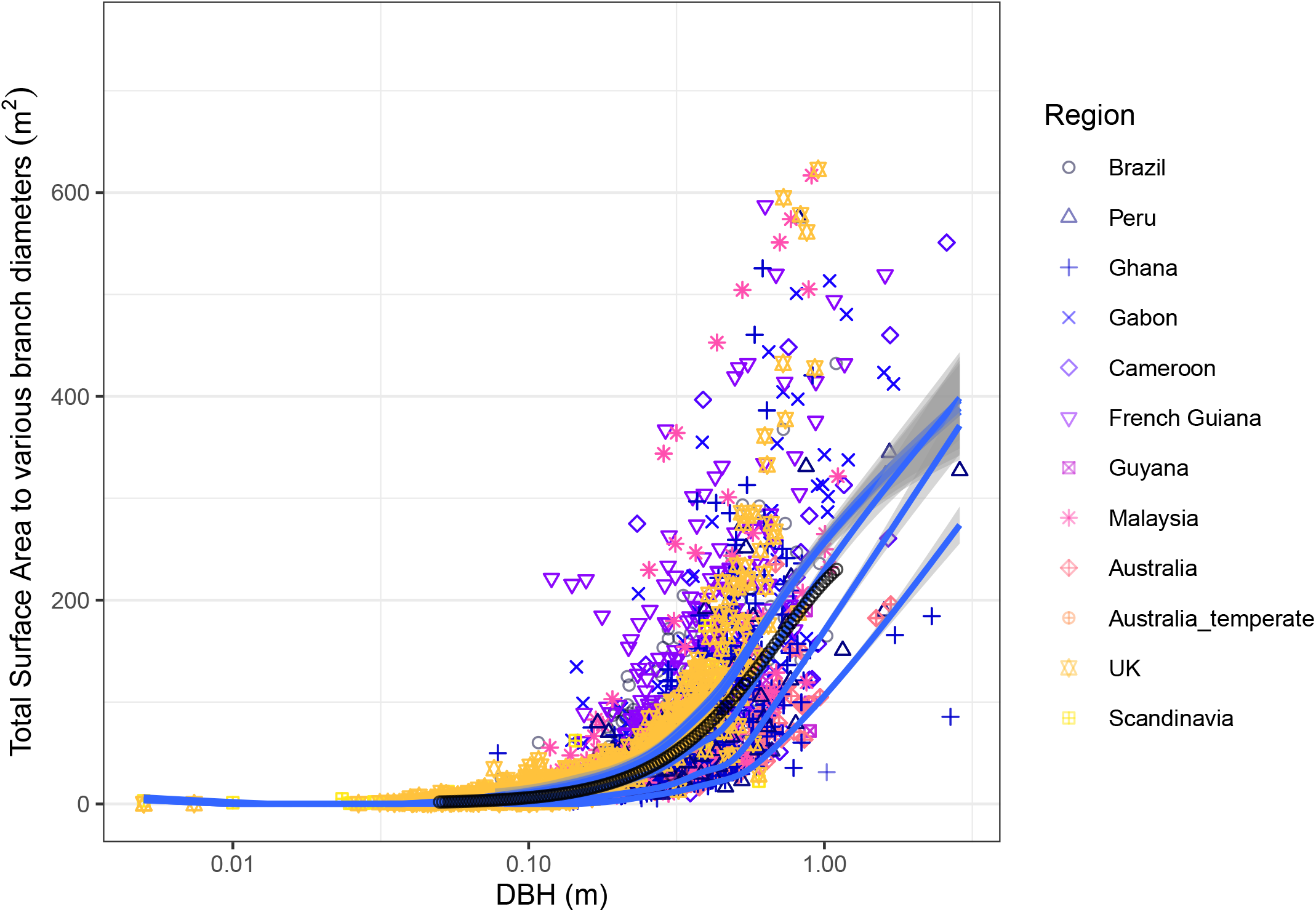
Same data as the figure above plotted on a linear scale. Each blue line includes increasingly fine branch diameters: 20cm (lowest line), 10cm, 5cm, 2cm, 1cm, and including all branches (top line). Black dots are expected values of Chambers (2004) allometry.

**Figure 7.**
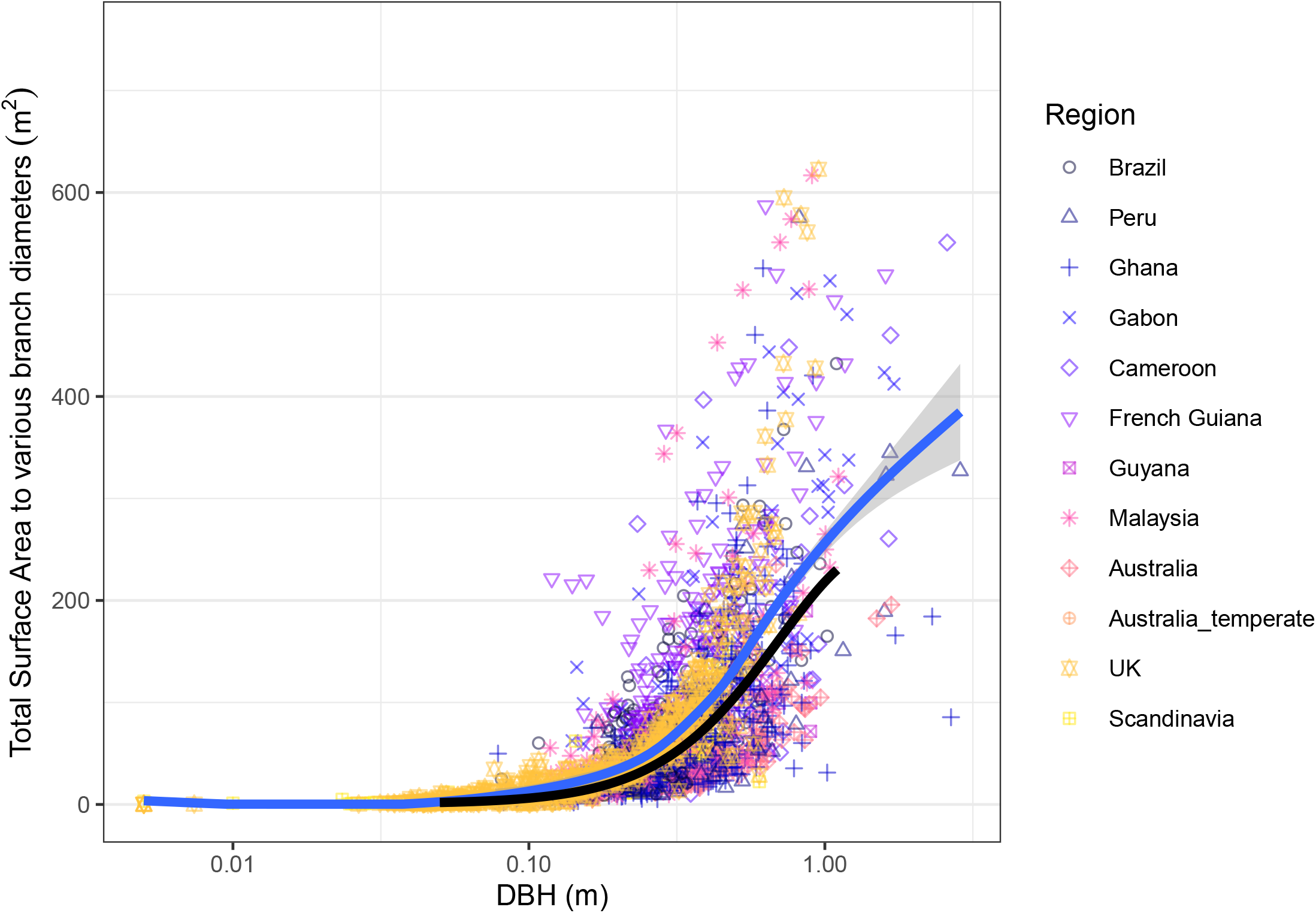
Same plot as above, but just for 0cm branches.

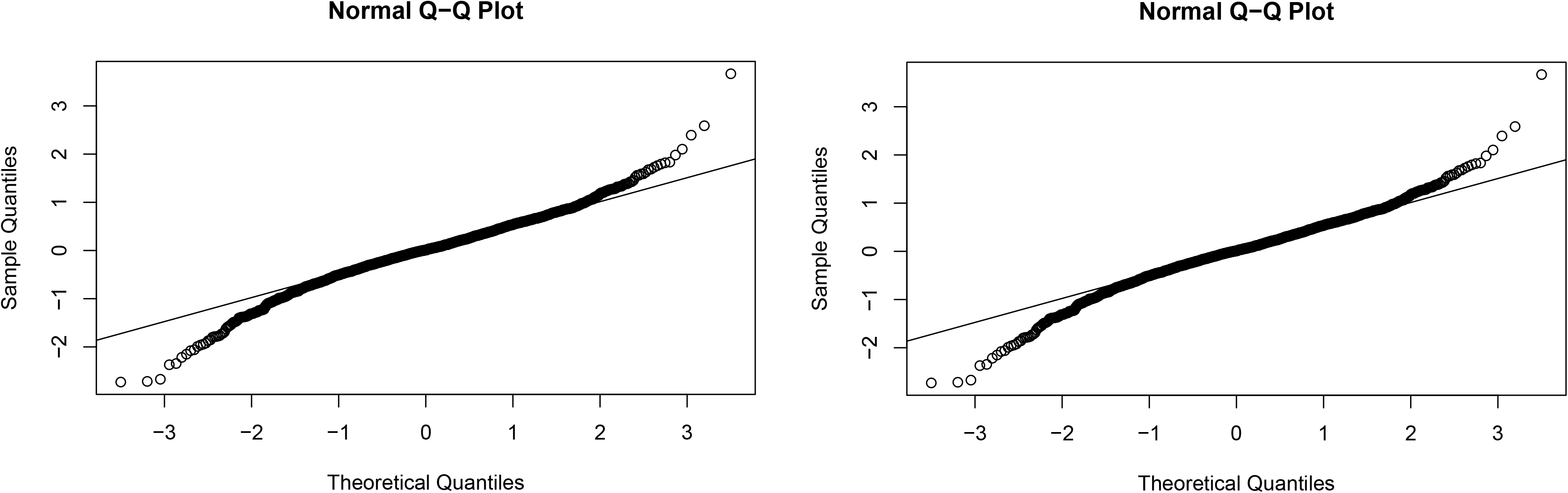

The linear model doesn’t seem to do well at the ends, which may indicate the the scaling parameter is changing with tree size. We need to fit non-linear models. There are a number of ways to do this. The *nlme* package is ok, but the community hasn’t figured out how to get confidence intervals and predictor significance out of these kinds of models, as far as I can tell. Another way is turning to Bayesian methods. The advantages there are that we can specify exact model structures and produce allometric equations for use by others. The downside is the potential difficulty in implementing these models; there are R packages to do this (e.g. rstanarm), but we might have to turn to STAN if we want specific things, and that’s a learning curve. Finally, there are GAMs. The upside is that they’re relatively simple and flexible. The downside is that it’s hard (impossible?) to constrain their functional forms, and I think it’s unlikely we’d be able to get allometric equations out of them for publication. Nonetheless, they should be good for data exploration, so let’s start there and see how it goes.

### GLMM

This is the basic GLMM we’re using: `glmer(surface_area_tot ∼ log(dbh) + (11plot_code), data = treestructs_all, family=Gamma(link=“log”))`

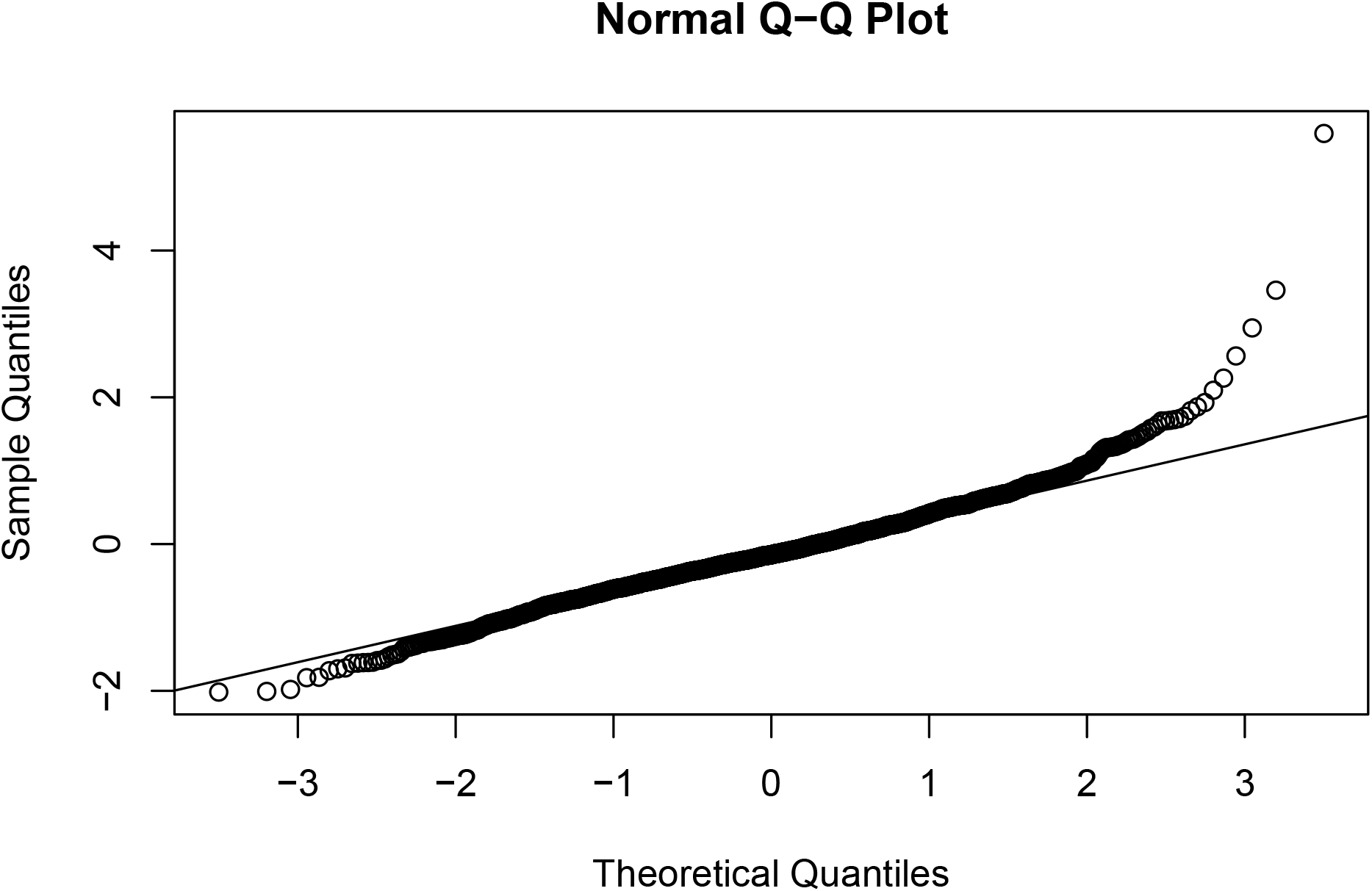

### Testing polynomial terms

Chambers (2004) employs polynomial terms in his log-log models. Including both powers of 2 and 3 improve the model, though those terms do not improve the model individually. We exclude those terms for now, and stick with the log(dbh) predictor only, especially given that the biological significance of power terms in a log-log model are unknown.

~~~
##
## Model selection based on AICc:
##
## K AICc Delta_AICc AICcWt Cum.Wt LL
## log(dbh) + log(dbh)^2 + log(dbh)-3 6 19732.60 0.00 1 1 -9860.28
## log(dbh) + log(dbh)^3 5 19797.14 64.55 0 1 -9893.56
## log(dbh) + log(dbh)^2 5 19822.64 90.04 0 1 -9906.30
## log(dbh) 4 19847.23 114.63 0 1 -9919.60
~~~

### GAM analysis

The models we’re using for GAM fits is:

**Figure 8.**
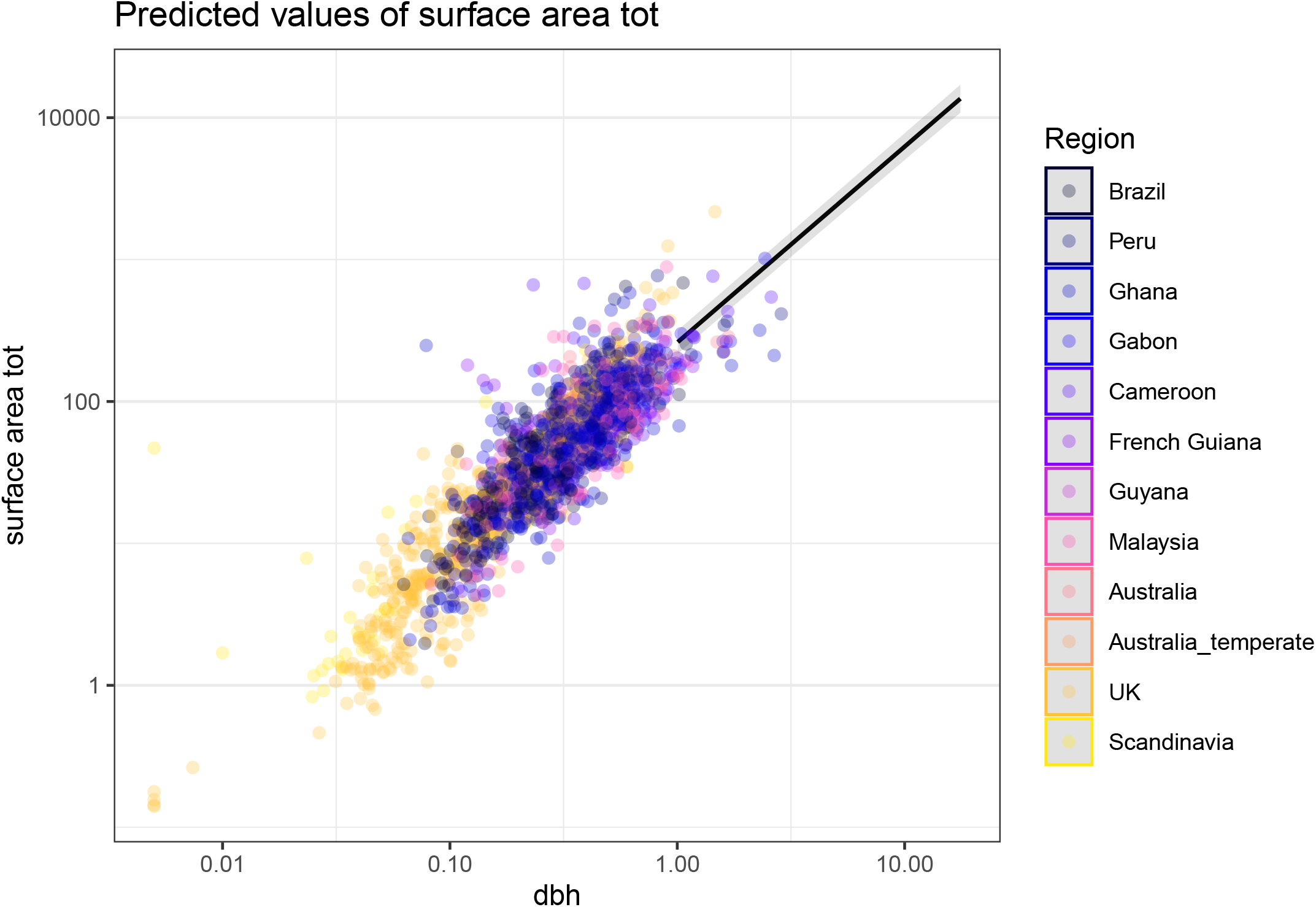
LMM marginal plot for surface area vs dbh. Points are residuals (i.e. plot effects have been removed).

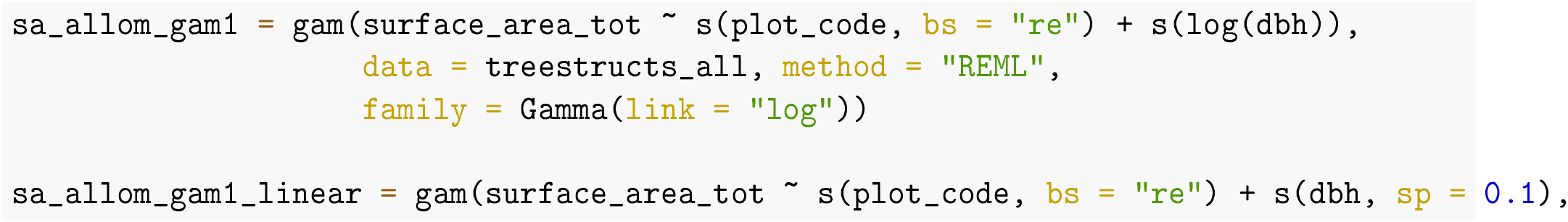
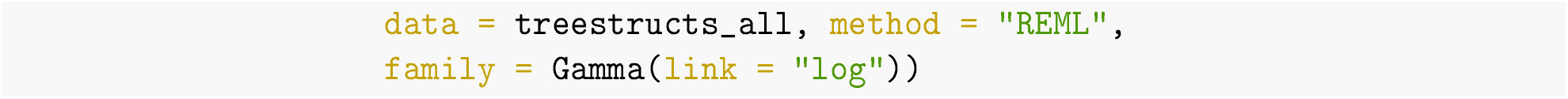

The log predictor is better than the linear (not surprising, delta AIC = 152.7463509). So we use the log formulation below.

Let’s look at that model, predicted across plots.

**Figure 9.**
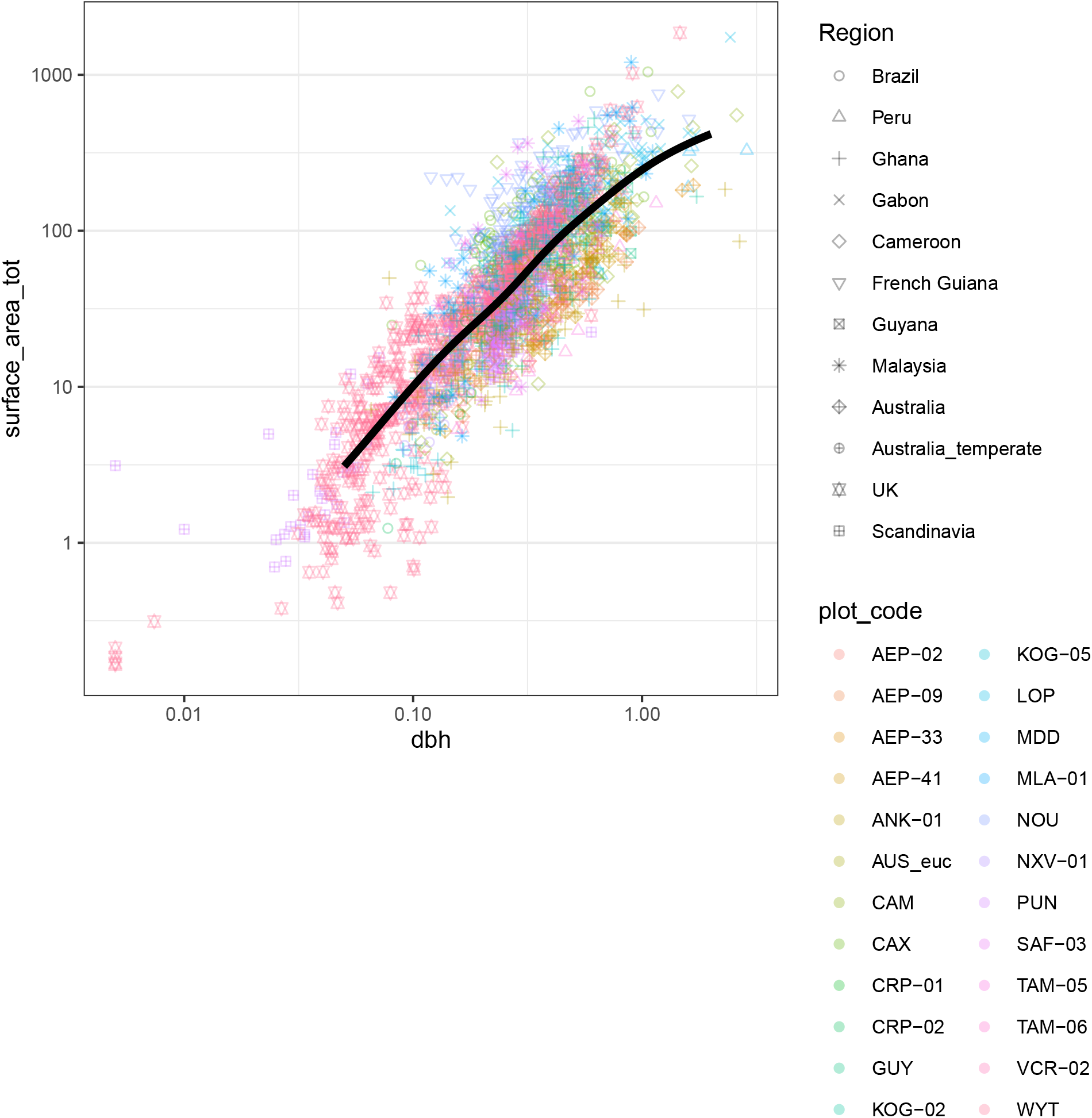
AMM fit across all plots

The fit does not go through the middle of the points due to the plot random effect weighting. The GAM looks good when looking at the predictions across each plot.

### HGAM analysis

Pedersen et al (2019) describe employing GAMs with a hierarchical structure (HGAMs) that allows shapes to change across groups in a penalized manner. We employ HGAMs here for the purposes of prediction. We do this because research into allometric relationships often seeks the best empirical fit of a model to d ata. These models thus often include polynomial predictors, for example, that have no biological interpretation. Such inclusion is warranted when one is seeking the best empirical fit p ossible. Nonetheless, it begs the question, if one is inlcluding the 1st, 2nd, and 3rd power in a model, why not also test the 1.2 power? Or the 4.6 power? That isto say, if we are seeking an empirical fit, we should be guided by the shape of the data and not our preconceived notions of polynomials or otherwise. GAMs can be a good tool to employ in this situation: they are flexible, penalizable, and mature.

**Figure 10.**
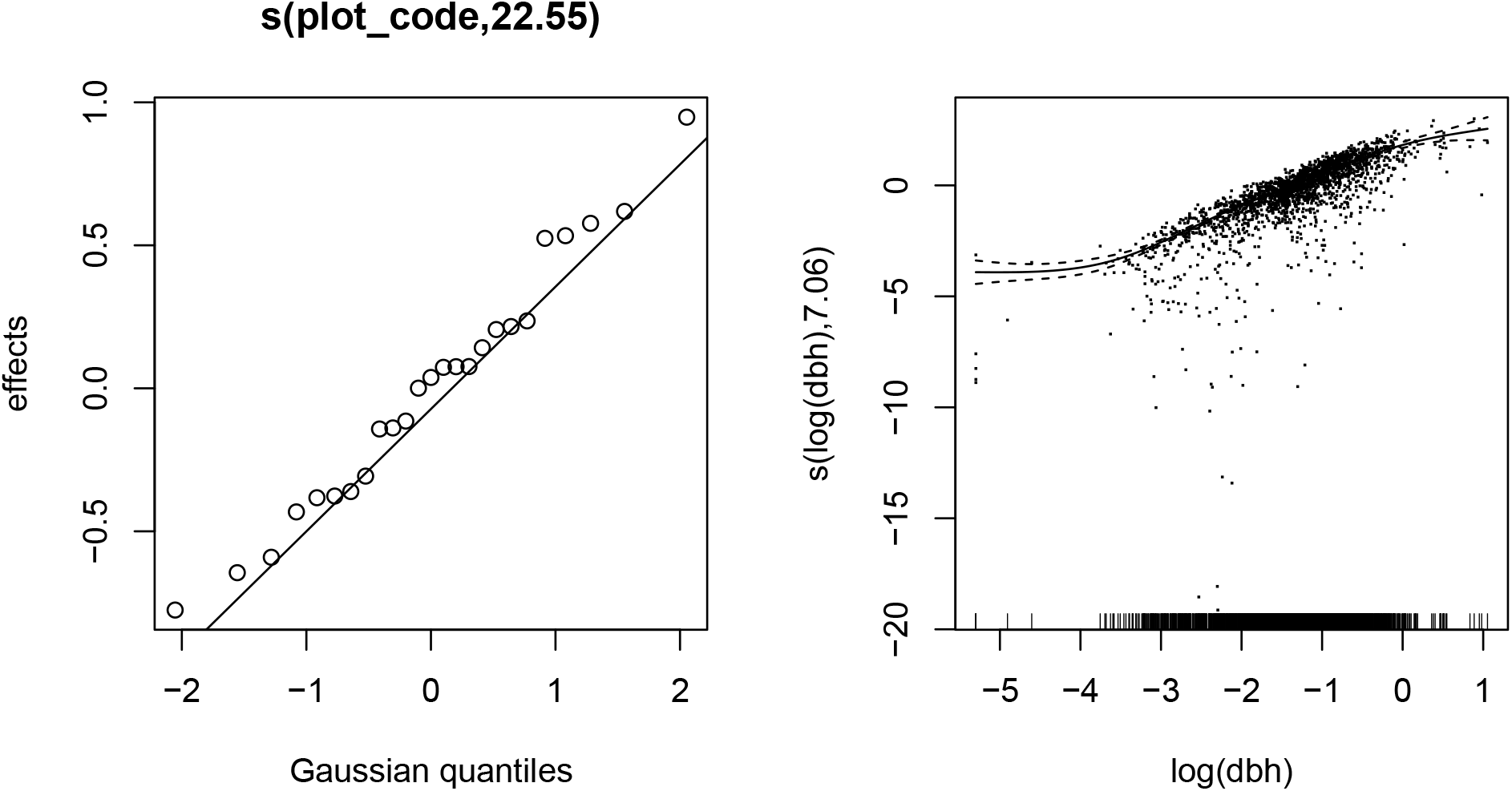
GAM checks

**Table 2:** AIC comparison of GAM models.

| mod | df | AIC | delta_AIC |
| --- | --- | --- | --- |
| hgam_eco_GI | 89.1 | 19327.62 | 0.0 |
| hgam_eco_GS | 100.4 | 19343.58 | 16.0 |
| hgam_GS | 65.4 | 19358.99 | 31.4 |
| hgam_eco_randint | 54.5 | 19432.26 | 104.6 |
| sa_allom_gam1 | 31.9 | 19681.97 | 354.4 |
| hgam_eco_norand_GI | 34.2 | 20271.28 | 943.7 |

The best model is

To test differences across ecosystems, we can look at differences between smooths per Gavin Simpson: https://www.fromthebottomoftheheap.net/2017/10/10/difference-splines-i/

### Breakpoint analysis

The GAM looks pretty ok as it is, but it’s hard to get equations out of a GAM that you can test theory with, and that you can produce equation with for an allomtry. So, let’s look into a breakpoint analysis to see if that low end residual problem can be solved. We can try this using the *segmented* package:

**Figure 11.**
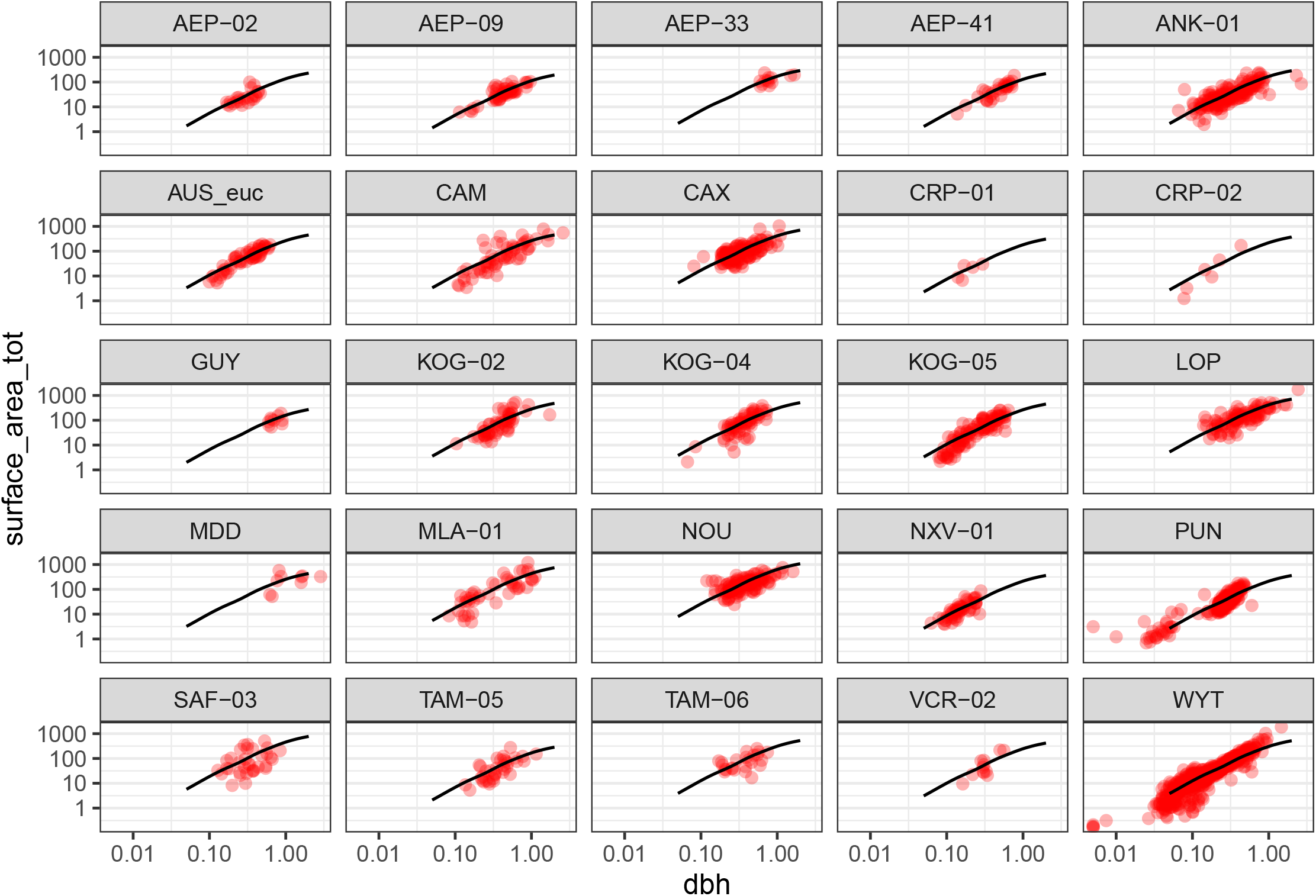
GAM allometry fits across plots

**Figure 12.**
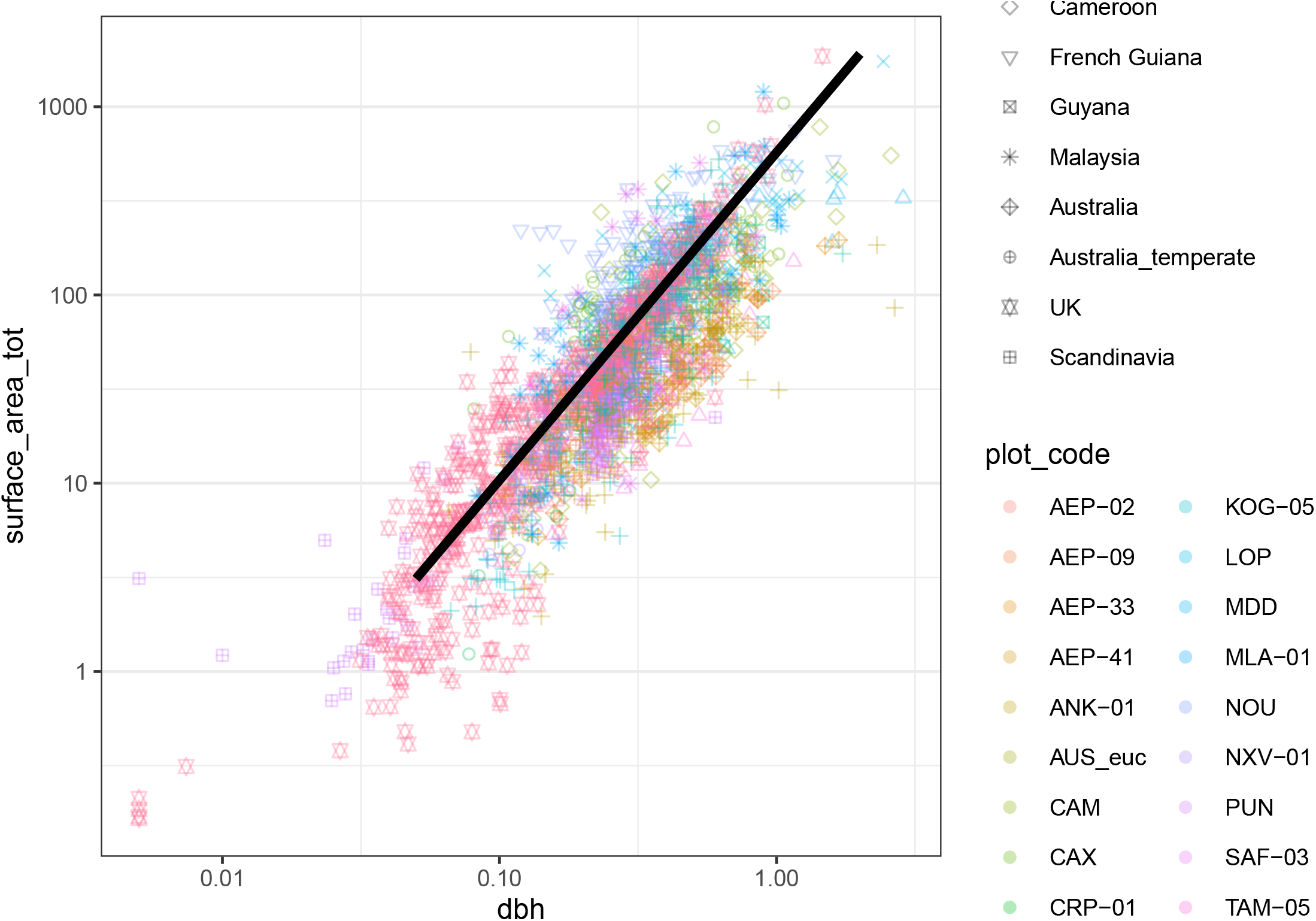
Global HGAM fit across all plots

~~~
##
## ***Regression Model with Segmented Relationship(s)***
##
## Call:
## segmented.lm(obj = sa_allom_lin, seg.Z = ∼dbh_log, psi = -1.6)
##
## Estimated Break-Point(s):
## Est. St.Err
## psi1.dbh_log -0.699 0.089
##
## Meaningful coefficients of the linear terms:
## Estimate Std. Error t value Pr(>1t1)
## (Intercept) 5.12117 0.12080 42.394 < 2e-16 ***
## dbh_log 1.58699 0.02379 66.720 < 2e-16 ***
## plot_codeAEP-09 -0.14809 0.14234 -1.040 0.298284
## plot_codeAEP-33 0.27718 0.20481 1.353 0.176090
## plot_codeAEP-41 -0.01946 0.15752 -0.124 0.901688
## plot_codeANK-01 0.18111 0.12566 1.441 0.149640
## plot_codeAUS_euc 0.74819 0.13796 5.423 6.52e-08 ***
## plot_codeCAM 0.48972 0.14020 3.493 0.000487 ***
## plot_codeCAX 1.11542 0.12671 8.803 < 2e-16 ***
## plot_codeCRP-01 0.28619 0.28720 0.996 0.319143
## plot_codeCRP-02 0.35460 0.26673 1.329 0.183837
## plot_codeGUY 0.19201 0.22248 0.863 0.388200
## plot_codeK0G-02 0.66227 0.14068 4.708 2.67e-06 ***
## plot_codeK0G-04 0.76602 0.13748 5.572 2.84e-08 ***
## plot_codeK0G-05 0.67389 0.12913 5.219 1.98e-07 ***
## plot_codeL0P 1.11540 0.13101 8.514 < 2e-16 ***
## plot_codeMDD 0.53191 0.23712 2.243 0.024988 *
## plot_codeMLA-01 1.12692 0.14661 7.687 2.29e-14 ***
## plot_codeN0U 1.52428 0.12663 12.037 < 2e-16 ***
## plot_codeNXV-01 0.57691 0.14072 4.100 4.29e-05 ***
## plot_codePUN 0.49334 0.12276 4.019 6.05e-05 ***
## plot_codeSAF-03 0.92325 0.15464 5.970 2.77e-09 ***
## plot_codeTAM-05 0.11843 0.15270 0.776 0.438083
## plot_codeTAM-06 0.78286 0.16771 4.668 3.23e-06 ***
## plot_codeVCR-02 0.55659 0.20589 2.703 0.006920 **
## plot_codeWYT 0.84066 0.12098 6.949 4.87e-12 ***
## U1.dbh_log -0.57445 0.10841 -5.299 NA
## ---
## Signif. codes: 0 ‘***’ 0.001 ‘**’ 0.01 ‘*’ 0.05 ‘.’ 0.1 ‘ ‘ 1
##
## Residual standard error: 0.5859 on 2133 degrees of freedom
## Multiple R-Squared: 0.8114, Adjusted R-squared: 0.809
##
## Convergence attained in 4 iter. (rel. change 6.4084e-07)
~~~

So, the segmented regression converges on the most likely breakpoint to be 50cm DBH when we start our search at 20cm. That seems reasonsable (even quite good), but *segmented* uses lm, and we want random effects. Let’s try that with basis functions.

**Figure 13.**
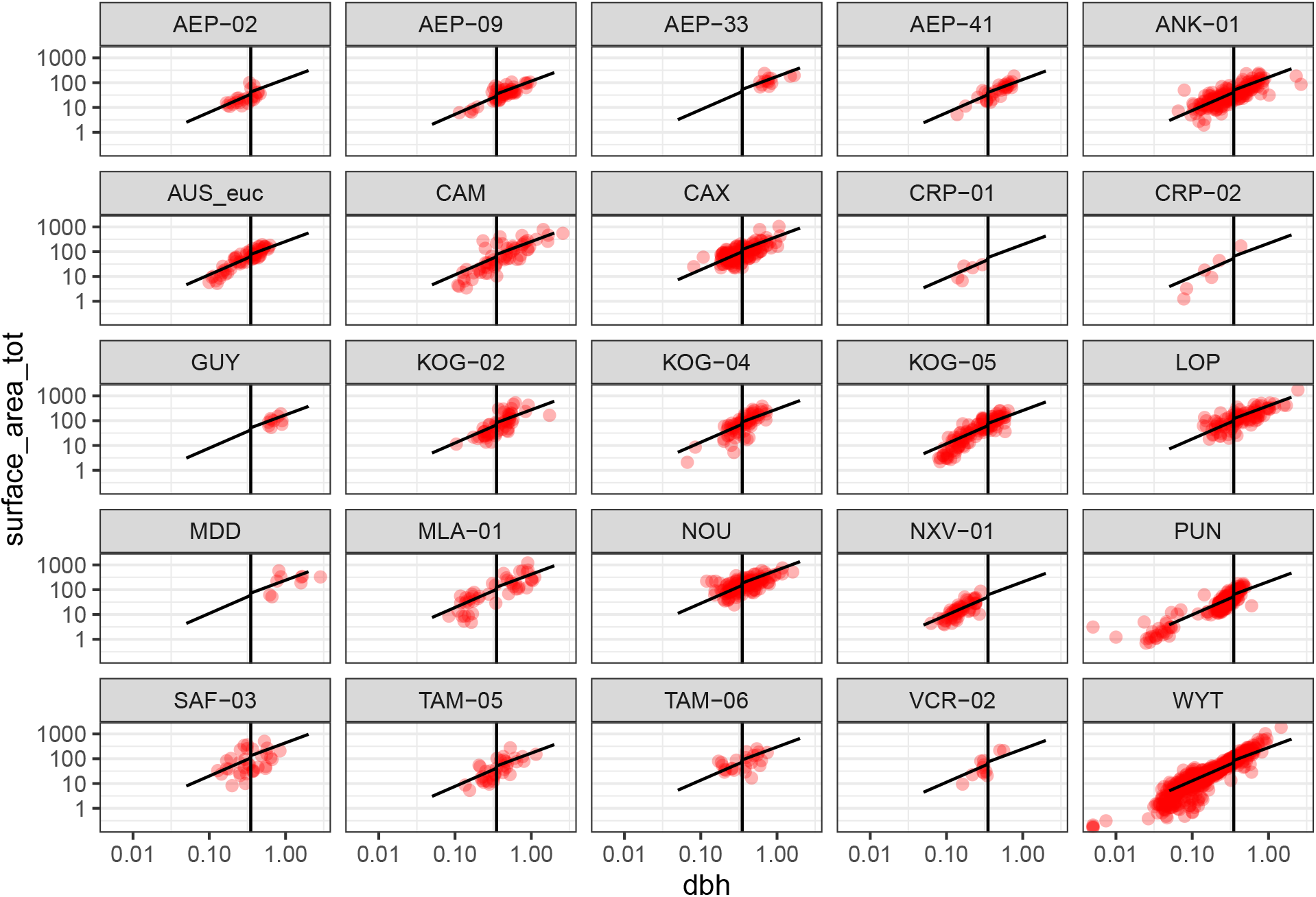
Piecewise GLMM

The piecewise GLMM finds the breakpoint at 35cm when we restrict the breakpoint search between 10 and 50cm. This is different from the linear model as it’s a completely different model, with random effect of plot and a gamma error structure. Nonetheless, the exponent difference is small.

We now fit one breakpoint between between 10 and 50cm (breakpoint fit = 35cm), and another between between 50 and 200cm (breakpoint fit = 87cm). We fit 3 models: 1 with a breakpoint at 35cm, another with a breakpoint at 87cm, and finally, another with both breakpoints. Comparing these models via AIC with the GLMM with no breakpoints results in the following:

~~~
##
## Model selection based on AICc:
##
## K AICc Delta_AICc AICcWt Cum.Wt LL
## Breakpoints at 35 and 87cm 6 19818.52 0.00 0.8 0.8 -9903.24
## Breakpoint at 35cm 5 19821.34 2.82 0.2 1.0 -9905.66
## Breakpoint at 87cm 5 19836.01 17.49 0.0 1.0 -9912.99
## No breakpoints 4 19847.23 28.70 0.0 1.0 -9919.60
~~~

That is, one breakpoint makes a big difference, and the 35cm breakpoint is significantly better than the 87cm breakpoint model, and while the model with both breakpoints does add some information to the model, it is only marginally better (in some cases, depending on which datasets we use). Thus, for simplicity, we stick with the 35cm breakpoint as our go-to model (for now).

### Comparing breakpoint and GAM models

Do GAMs do a better job of modeling surface area allometry than breakpoint models? We should probably use gamm4 instead of mgcv when comparing gam’s with glmm’s, as gamm4 uses lme4, and it (I think) uses similar logic for determining the degrees of freedom of random effects as does lme4. Hence, comparing gamm4 vs lme4 models should be more reliable. One difference is that gamm4 doesn’t support spline penalties, which was important to avoid over-fitting.

The AIC of the gamm4 model is better than the one breakpoint model (AIC(gamm4 - break- point) = -84.31). Note that K (i.e. parameter count) of the gamm4 model (K = 5) is reported to be the same as the breakpoint model. This is an outstanding question in stats, it seems… That is, how to compare GAMMs and GLMMs is unresolved… (see https://stat.ethz.ch/pipermail/r-help/2010-January/225829.html)

### Testing scaling theory

The scaling coefficients resulting from the breakpoint model are the following: alpha = 1.34 for trees up to 35cm DBH, and alpha = 1.14 for trees larger than 35cm DBH.

### Surface area across branch size classes

**Figure 14.**
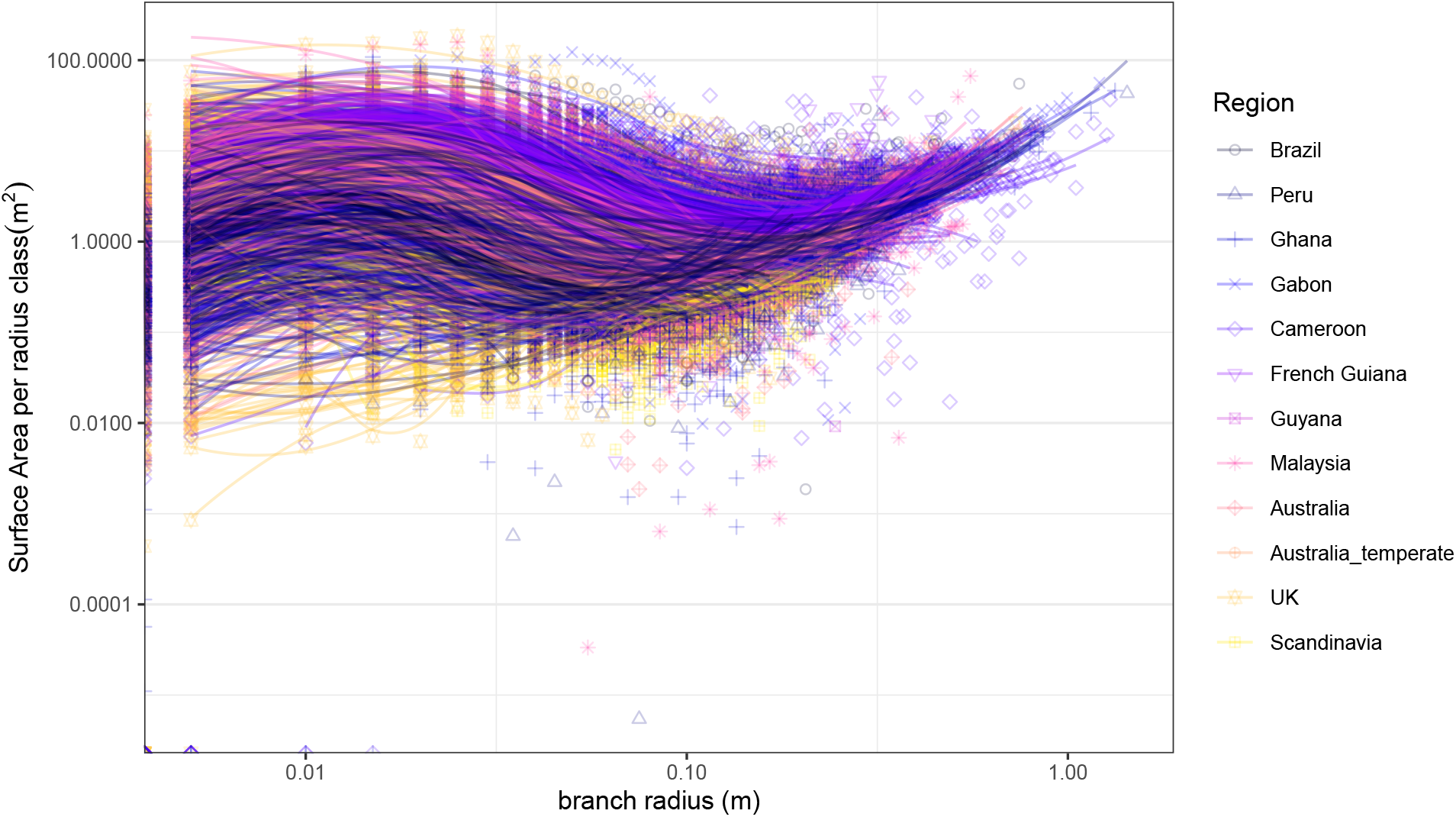
Surface area across branch size classes, lines per tree

**Figure 15.**
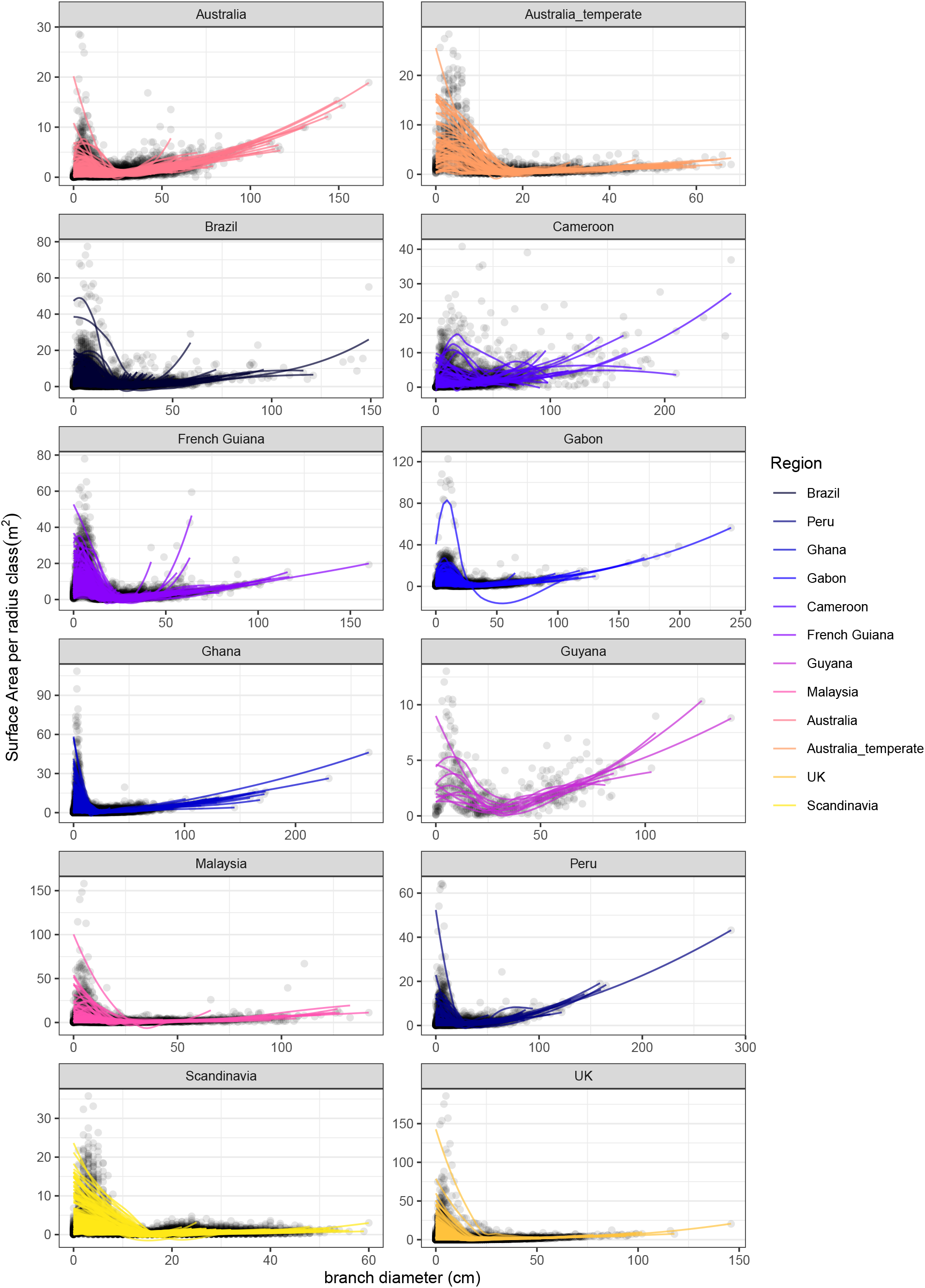
Surface area across branch size classes, lines per tree 29

**Figure 16.**
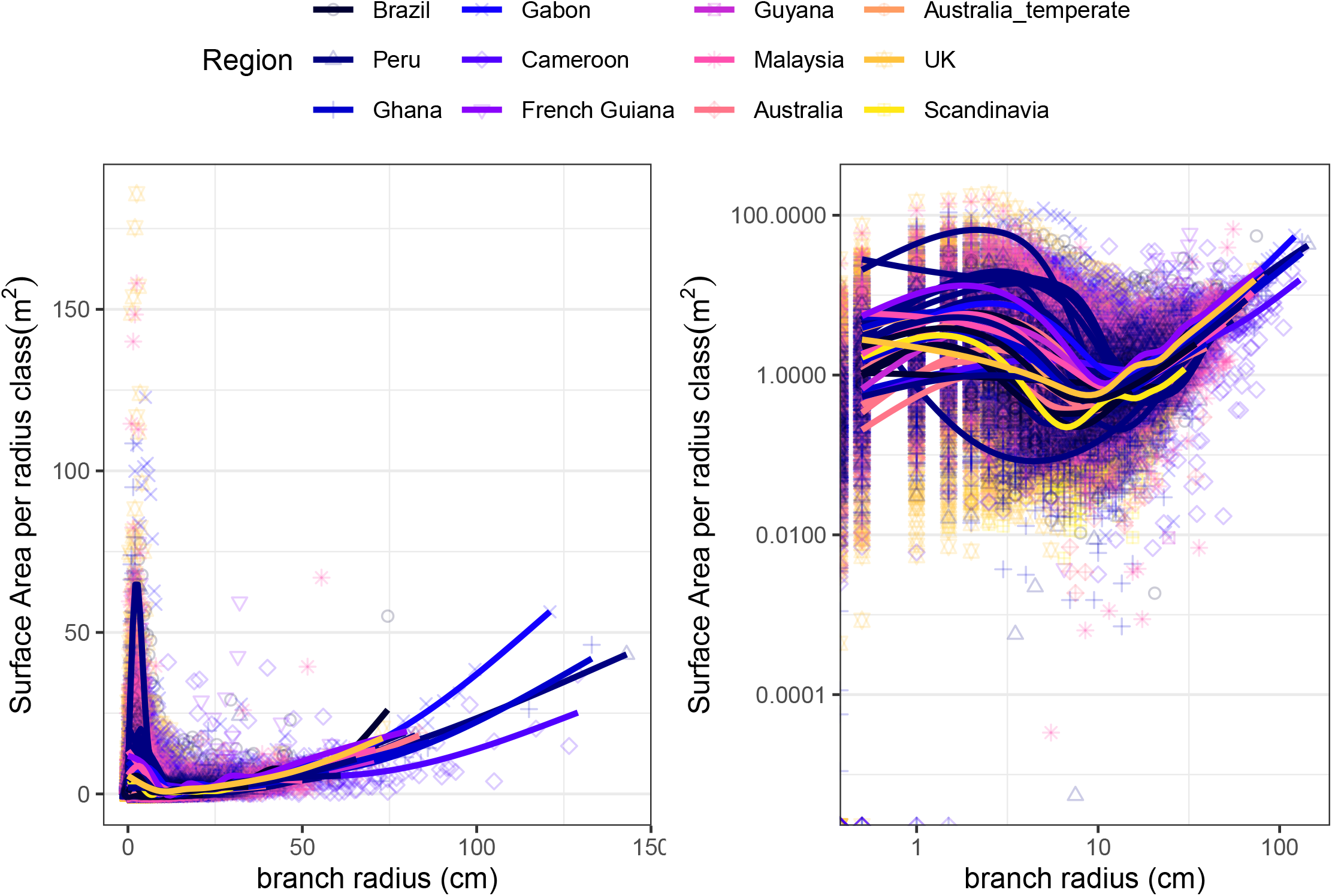
Surface area across branch size classes, lines per plot

### Surface area across branch size classes

Since surface area ∝ *r*, and volume (biomass) ∝ *r*^2^, volume decreases more quickly with branch size than does surface area. Thus, we would expect smaller branches to harbor a larger proportion of a tree’s surface area than a tree’s biomass.

**Figure 17.**
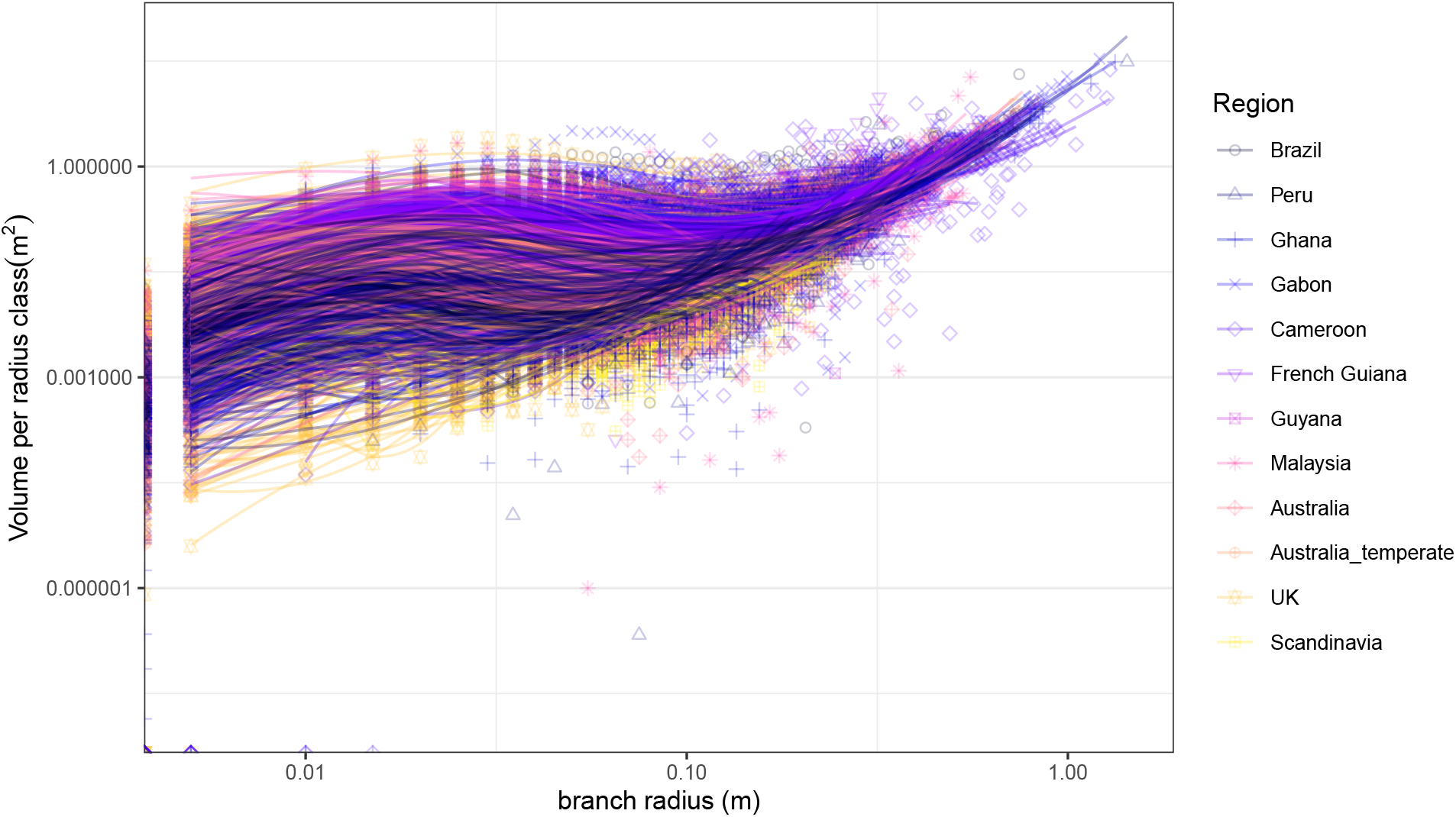
Volume across branch size classes, lines per tree

